# Regulated assembly and neurosteroid modulation constrain GABA_A_ receptor pharmacology *in vivo*

**DOI:** 10.1101/2023.02.16.528867

**Authors:** Chang Sun, Hongtao Zhu, Sarah Clark, Eric Gouaux

**Affiliations:** Vollum Institute, Oregon Health and Science University, 3232 SW Research Drive, Portland, Oregon 97239; Howard Hughes Medical Institute, Oregon Health and Science University, 3232 SW Research Drive, Portland, Oregon 97239

## Abstract

Type A GABA receptors (GABA_A_Rs) are the principal inhibitory receptors in the brain and the target of a wide range of clinical agents, including anesthetics, sedatives, hypnotics, and antidepressants. However, our understanding of GABA_A_R pharmacology has been hindered by the vast number of pentameric assemblies that can be derived from a total 19 different subunits and the lack of structural knowledge of clinically relevant receptors. Here, we isolate native murine GABA_A_R assemblies containing the widely expressed α_1_ subunit, and elucidate their structures in complex with drugs used to treat insomnia (zolpidem and flurazepam) and postpartum depression (the neurosteroid allopregnanolone). Using cryo-EM analysis and single-molecule photobleaching experiments, we uncover only three structural populations in the brain: the canonical α_1_β2γ_2_ receptor containing two α_1_ subunits and two unanticipated assemblies containing one α_1_ and either an α_2_, α_3_ or α_5_ subunit. Both of the noncanonical assemblies feature a more compact arrangement between the transmembrane and extracellular domains. Interestingly, allopregnanolone is bound at the transmembrane α/β subunit interface, even when not added to the sample, revealing an important role for endogenous neurosteroids in modulating native GABA_A_Rs. Together with structurally engaged lipids, neurosteroids produce global conformational changes throughout the receptor that modify both the pore diameter and binding environments for GABA and insomnia medications. Together, our data reveal that GABA_A_R assembly is a strictly regulated process that yields a small number of structurally distinct complexes, defining a structural landscape from which subtype-specific drugs can be developed.

## Main

Regulation of brain excitability by activation of neuronal GABA_A_Rs is essential for normal brain development and function^1–3^. Deficits in GABA_A_R activity are associated with health problems ranging from epilepsy to intellectual disability^4^. A large number of ions and small molecules modulate GABA_A_R activity, including Zn^2+^ (ref. 5) and picrotoxin^6^ as well as therapeutic agents such as benzodiazepines^6–8^, barbiturates^8^, and propofol^8^. Indeed, the GABA_A_R modulators flurazepam and zolpidem are widely used to treat insomnia. Lipophilic neurosteroids are also potent endogenous modulators of GABA_A_Rs. Allopregnanolone (ALP; 3α-OH-5α-pregnan-20-one), synthesized chiefly in the brain^9–11^, potentiates GABA_A_R activity in a subunit-dependent manner^12–14^, and its anxiolytic and sedative effects have proved to be effective for the treatment of postpartum depression^15^. In addition, ganaxolone, a synthetic mimetic of allopregnanolone, has recently entered the clinic as an anticonvulsive agent.

Because modulation of receptor function is dependent upon subunit composition and arrangement, a knowledge of native GABA_A_R architecture is crucial to understand how these different molecules elicit distinct physiological responses. However, the potential diversity of pentameric GABA_A_Rs is vast due to the existence of 19 different receptor subunits (α_1-6_, β_1-3_, γ_1-3_, ρ_1-3_, δ, ε, π, and θ). Moreover, studies *in vitro* suggest that variations in subunit expression levels can modify subunit stoichiometry. Despite progress in resolving the architecture of recombinant di- and tri-heteromeric GABAARs^5, 7, 16–18^, there is no structural understanding of the various GABA_A_Rs that are present in the brain. Indeed, although the presence of specific subunits in native receptor assemblies has been determined by immunoprecipitation^19–21^, the number and arrangement of subunits remains unknown.

To elucidate the ensemble of GABA_A_Rs that define the molecular action of endogenous and therapeutic modulators, we isolated native *α*_1_ subunit-containing GABA_A_Rs (n*α*_1_GABA_A_Rs) from mouse brain using an engineered high-affinity, subunit-specific Fab fragment^7^. Because the α_1_ subunit is ubiquitously expressed throughout the brain, and is a subunit of both synaptic and extrasynaptic receptors, this approach enabled us to analyze 60%–80% of native GABA_A_Rs ^22–24^. Furthermore, it permitted an investigation of three clinically-relevant molecules, allopregnanolone (ALP), didesethylflurazepam (DID), and zolpidem (ZOL), all of which target *α*_1_-containing receptors. With isolated n*α*_1_GABA_A_Rs in hand, we were able to count the number of α_1_ subunits in these complexes by single-molecule fluorescence bleaching experiments, investigate protein composition by mass spectrometry, and elucidate high resolution structures of n*α*_1_GABA_A_R assemblies by single-particle cryo-EM. Our results reveal a surprisingly small number of receptor complexes, whose structures provide a framework from which targeted drugs could be developed.

### Isolation of functional n*α*_1_GABA_A_Rs from the mouse brain

We engineered the 8E3 *α*_1_ subunit-specific Fab fragment^7^ to include a GFP fluorophore, affinity tag, and 3C-protease site to enable release it the affinity resin (8E3-GFP; K_d_ = 0.5 nM; **Extended Data Figure 1**). The 8E3-GFP Fab was then used to isolate n*α*_1_GABA_A_Rs from solubilized mouse brain tissue (excluding the cerebellum) while monitoring the purification workflow via GFP fluorescence. Detergent (lauryl maltose meopentyl glycol) treatment routinely solubilized the majority of n*α*_1_GABA_A_Rs, accompanied by the inhibitory synapse marker neuroligin2 (**Extended Data Figure 1**). Following nearly complete capture of receptors on affinity resin (**Extended Data Figure 1**), n*α*_1_GABA_A_Rs complexes were reconstituted into lipid-filled nanodiscs^25^ then eluted by 3C-protease treatment. Further purification by size exclusion chromatography yielded an ensemble of n*α*_1_GABA_A_R:Fab complexes. Radioligand binding assays showed that the purified pentameric preparations were functional and retained high-affinity flunitrazepam binding (K_d_ = 6.0 ± 0.2 nM (mean ± s.e.m.); **Extended Data Figure 1**)^26^. Furthermore, analysis of the purified native receptor complexes by mass spectrometry identified all α and β subunits as well as the γ_1_, γ_2_, and δ subunits, demonstrating that α_1_-dependent isolation captured receptors containing most of the 19 GABA_A_R subunits.

### n*α*_1_GABA_A_Rs comprise three structural populations

To elucidate the composition and arrangement of native receptors, we collected cryo-EM data from n*α*_1_GABA_A_R:Fab complexes in the presence of DID, ZOL plus GABA, and ALP plus GABA (**Extended Data Table 1**), and carried out single particle analysis. The 2D class averages derived from all three datasets showed prominent Fab features at the periphery of the receptors. In contrast to a previous study on recombinant *α*_1_-containing tri-heteromeric GABA_A_R complexes in which all receptors contained two *α*_1_ subunits^7^, we observed class averages with only one Fab bound (**Extended Data Figure 2–4**), demonstrating the presence of receptors with a single α_1_ subunit.

We subsequently used extensive 3D classification to rigorously define subunit composition and arrangement of n*α*_1_GABA_A_R:Fab complexes. An inverse mask of the entire transmembrane domain (TMD) allowed us to exclude structural heterogeneity in the region of the pore and enabled classification to be driven by the *α*_1_-specific Fab and *N*-glycosylation patterns unique to each α, β, and γ subunit. After combining classes with the same Fab and *N*-glycosylation features, we consistently obtained three different 3D classes: a single class with two Fabs (two-Fab) and two classes with one Fab (one-Fab) (**Extended Data Figure 2–4**). We defined the two one-Fab classes as meta-one-Fab and ortho-one-Fab according to the relative position of their α_1_ and γ subunits. In all three classes from all three data sets we observed two α subunits, two β subunits, and one γ subunit arranged in an α*-β-α-β*-γ clockwise order when viewed from the extracellular side of the membrane (asterisks denote subunits adjacent to the γ subunit). This pentameric configuration therefore represents the dominant form of n*α*_1_GABA_A_Rs. Remarkably, we found no evidence for receptors with a β-β interface despite the high abundance of β subunits in native receptor assemblies, in contrast to recombinant α_1_-β_3_-α_1_-β_3_-β_3_^5^ and δ-containing^18^ assemblies. Thus, heterologous expression of GABA_A_ subunits appears to yield receptors in configurations that are not abundant in native brain tissue.

In the ALP/GABA dataset, 3D reconstructions at resolutions of 2.5 Å, 2.6 Å, and 2.6 Å were achieved for the two-Fab, ortho-one-Fab, and meta-one-Fab assemblies, respectively. This resolution was sufficient for subunit identification, small molecule positioning, and model building (**Figure 1a**; **Extended Data Figures 5–6; Extended Data Table 2**). The identities of β and γ subunits were determined from a combination of glycosylation patterns and sidechain densities. Receptors in the two-Fab class had a tri-heteromeric α_1_*-β_2_-α_1_-β_2_*-γ_2_ arrangement, consistent with genetic^27^, immunohistochemistry^21^, and electrophysiology^28^ data suggesting that this is the most abundant subtype in the brain.

**Figure 1.**
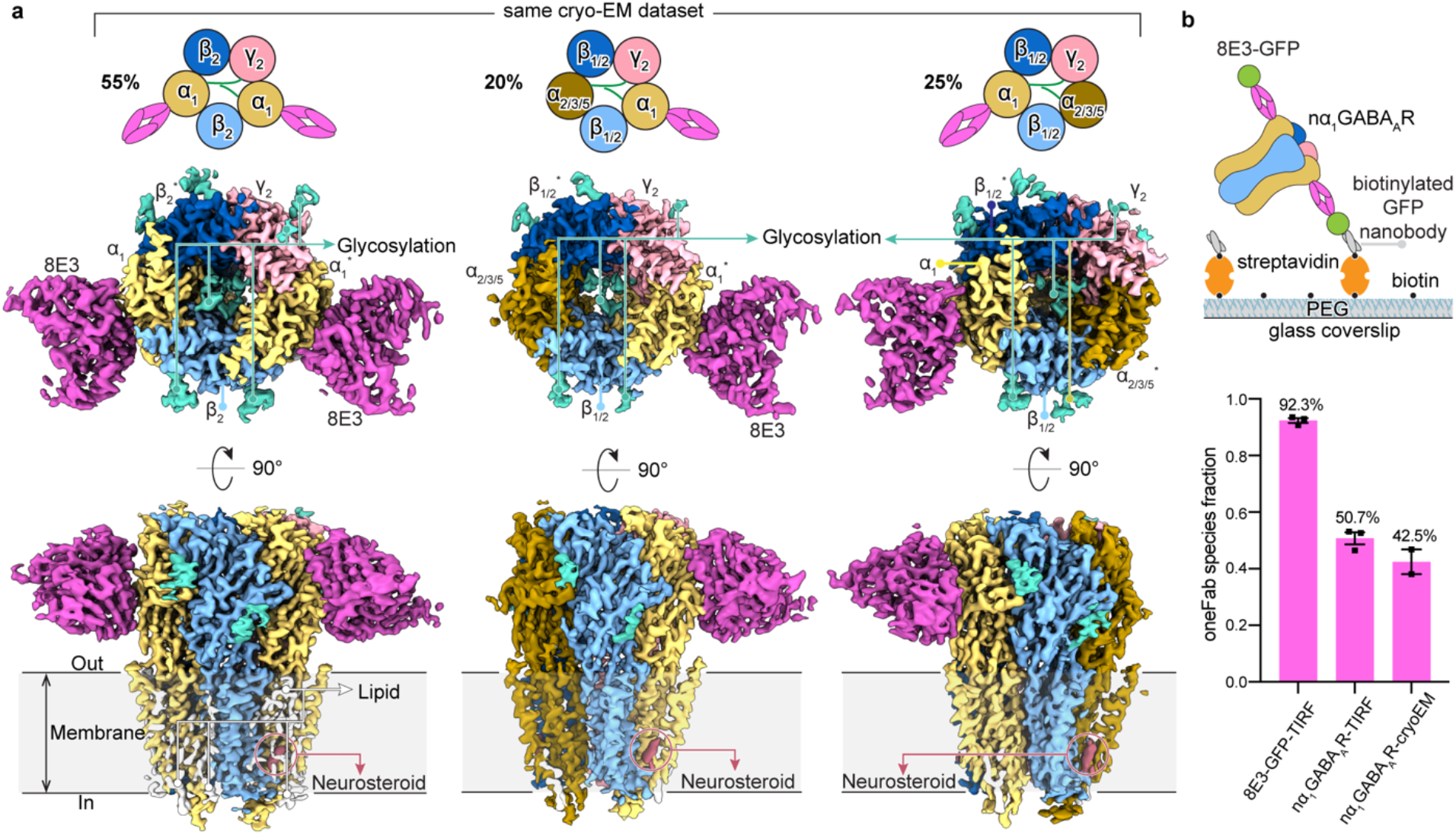
The three major nα_1_GABA_A_R complexes. **a,** Cryo-EM reconstruction of two-Fab α_1_*-β_2_-α_1_-β_2_*-γ_2_, ortho-one-Fab α_1_*-β_1/2_-α_2/3/5_-β_1/2_*-γ_2_ and meta-one-Fab α_2/3/5_*-β_1/2_-α_1_-β_1/2_*-γ_2_ receptor complexes (from left to right), purified from mouse brain using an α1-specific Fab. All reconstructions processed from the ALP/GABA dataset. Percentages calculated from particles separated by 3D classification using an inverse-TMD mask (Extended Data Figure 4). **b**, Single-molecule TIRF photobleaching of purified nα_1_GABA_A_R-GFP-Fab complexes. Top, experimental design; bottom, distribution of photobleaching events for nα_1_GABA_A_R-GFP-Fab complexes and isolated GFP-Fab (control). Individual data points are presented as squares while standard errors of the mean (SEM) are shown as error bars.

In contrast, each of the meta-one-Fab and ortho-one-Fab classes contained mixed receptor ensembles that we categorized as α_2/3/5_*-β_1/2_-α_1_-β_1/2_*-γ_2_ and α_1_*-β_1/2_-α_2/3/5_-β_1/2_*-γ_2_, respectively. A single α_1_ subunit in one position and either α_2_, α_3_, or α _5_ at the second *α* position, together with either *β*_1_ or *β*_2_ at the *β* position, yielded receptors with at least four and as many as five unique subunits in the pentameric assembly. Despite subunit ambiguity in the density maps, we evaluated the overall agreement between density map and protein sequences and modeled the α_2/3/5_ subunit from one-Fab structures as α_3_ and *β*_1/2_ as *β*_2_ to facilitate structural comparison among datasets. The common presence of a γ_2_ subunit in α_1_-containing GABA_A_Rs suggests a favorable association between these two highly expressed subunits. Furthermore, the highly ordered *N*-glycosylation of α subunits observed in the extracellular vestibule of the two-Fab class^7^ is conserved in both one-Fab classes, and includes a polysaccharide bridge between the γ_2_ subunit and the non-adjacent α subunit. Intriguingly, α_2/3/5_ subunits may have a fucose sugar attached to the asparagine-linked *N*-acetylglucosamine, which is absent in the α_1_ subunit (**Extended Data Figure 6**).

The two one-Fab classes comprise 45% of particles in the ALP/GABA dataset and 38% particles in the ZOL/GABA dataset (**Figure 1**; **Extended Data Figure 2–4**), demonstrating that receptors containing only one *α*_1_ subunit are more abundant than previously thought^29–31^. To independently measure the α_1_ subunit stoichiometry within n*α*_1_GABA_A_Rs, we measured photobleaching of the GFP fluorophore in purified 8E3-GFP complexes using single-molecule total internal reflection fluorescence (TIRF) microscopy^32^. Roughly 50% of photobleaching events comprised a single step (**Figure 1b**), indicating that about half the purified receptors have just one α_1_ subunit, in agreement with our cryo-EM data.

We used the two-Fab and ortho-one-Fab structures from the ALP/GABA dataset as paradigms to compare interdomain arrangements in receptors containing one or two α_1_ subunits. Despite containing highly homologous subunits (74% sequence similarity between α_1_ and α_3_), we observed striking differences between the one-Fab and two-Fab complexes. The extracellular domains (ECD)s and TMDs are almost identical in α_1_ and α_3_ subunits in equivalent positions, having backbone RMSDs of 0.45 Å and 0.35 Å, respectively. However, when aligned by TMD, the RMSD of ECD increases to 1.05 Å, suggesting significant inter-domain displacement between these α_1_ and α_3_ subunits. Furthermore, both meta-one-Fab and ortho-one-Fab have markedly shorter separations between the ECD and TMD center of masses (50.1/50.6 Å for meta-one-Fab α_3_*/α_1_ and 50.9/51.0 Å for ortho-one-Fab α_1_*/α_3_) than two-Fab complexes (51.9/52.0 Å for α_1_*/α_1_ subunits). This observed shortening is also apparent as a reduction of angles between the primary axes of the ECDs and the TMDs in one-Fab receptors (**Extended Data Figure 7**).

### ALP is a ubiquitous modulator of n*α*_1_GABA_A_Rs

Neurosteroids, such as ALP and allotetrahydrodeoxycorticosterone (THDOC), are endogenous ligands that confer anxiolytic, sedative, hypnotic, and anesthetic properties by potently and selectively potentiating GABA_A_Rs, and by direct activation at higher concentrations (≥ 100 nM)^33–36^. To investigate the molecular basis of neurosteroid modulation, we compared the structures of two-Fab, meta-one-Fab and ortho-one-Fab assemblies in complex with GABA and ALP (**Figure 2a**). In the two-Fab structure, two ALP molecules are bound in the TMD region, each approximately 60 Å ‘below’ one of the two GABA binding pockets in the ECD (**Figure 2b**). The ALP pockets are at the interface between transmembrane helices 1 and 4 (TM1 and TM4) of an α_1_ subunit and TM3 of the adjacent β_2_ subunit, which form an almost rectangular box lined by primarily aromatic and hydrophobic residues: α_1_-W245 on one side, β_2_-Y304 and β_2_-L301 at the base, and β_2_-L297, α_1_-V242 and α_1_-I238 on another side. Remarkably, lipid acyl chains are present on the other two long sides and the box is capped by α_1_-P400 and α_1_-Q241, the amide oxygen of the latter forming a hydrogen bond with the 3’-OH of ALP (**Figure 2c**).

**Figure 2.**
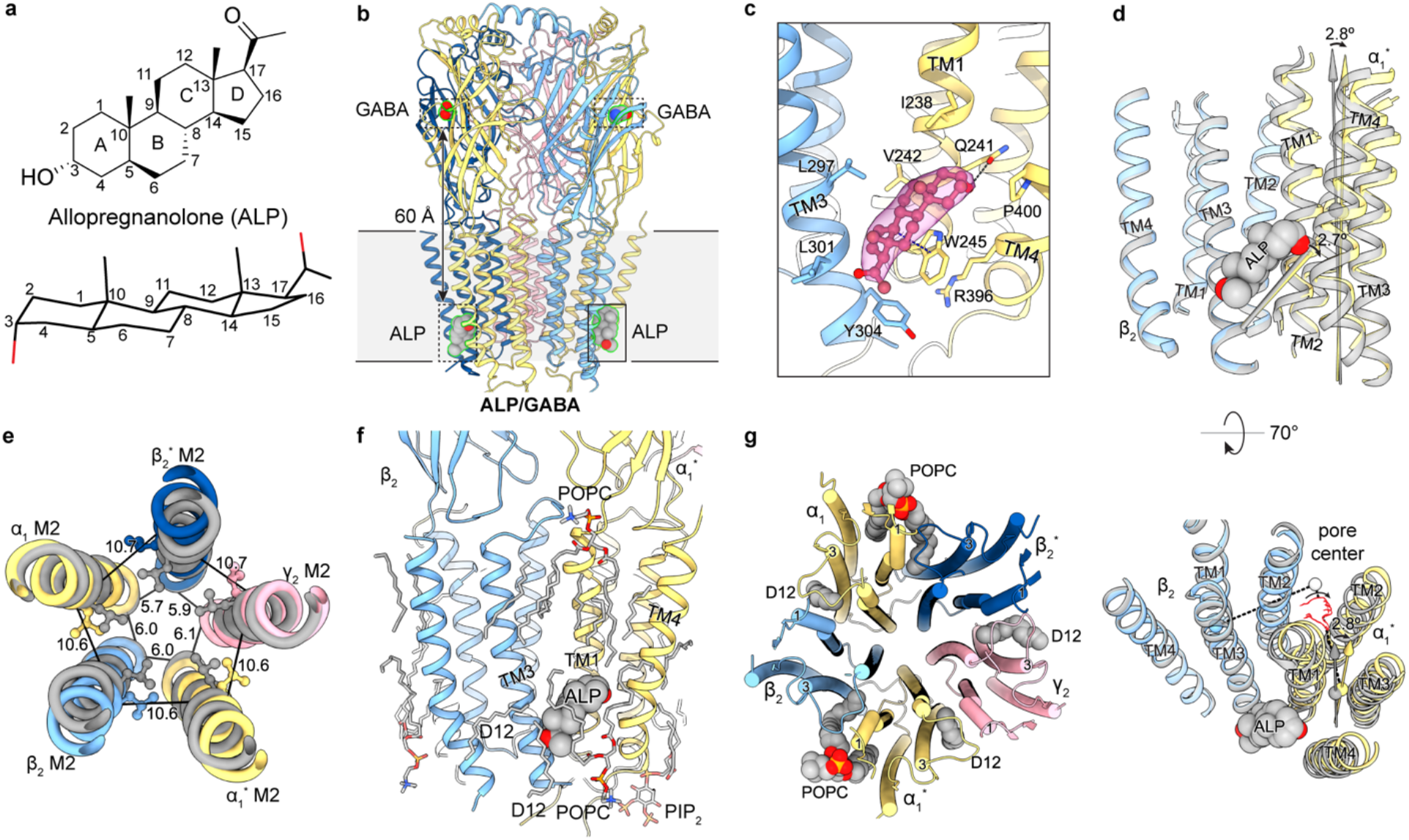
ALP sculpts the conformation of the TMD. **a**, Chemical structure of ALP. **b**, Structural overview of α_1_*-β_2_-α_1_-β_2_*-γ_2_ nα_1_GABA_A_R in complex with ALP and GABA. Bound Fabs hidden for clarity. **c**, Binding pose of ALP and ligand density in the binding pocket at the β_2_^+^/α_1_^*-^ TMD interface. **d**, Local conformational changes induced by ALP binding. Coordinates of gray structure of a full-length α_1_β_3_γ_2_ recombinant receptor in complex with GABA from PDB 6I53. Structural alignment based on the β subunit TMD. **e**, Global structural rearrangements induced by ALP binding. Structural alignment based on global TMD. Distances are between 9’ gate Cα and the -2’ gate. **f**, Lipids molecules resolved in the ALP/GABA structure. **g**, Lipids molecules with acyl tails inserted between TM1 and TM3 of adjacent subunits.

Incorporation of ALP remodels the conformation of TMD helices, enlarging the channel’s pore compared to a recombinant α_1_β3γ_2_ structure without neurosteroid (PDB 6I53). Local alignment of the β_2_ and α_1_* TMDs in the two structures reveals a 2.7° rotation of the line connecting the Cα atoms at the base and top of the ALP box (β_2_-Y304 and α*_1_-Q241) while the length of this line remains constant. In addition, the α_1_* TMD rotates by 2.8° around an axis between the center of mass of the entire TMD and the center of mass of the α*_1_ TMD (**Figure 2d**). We observed a similar but smaller effect at the β_2_*/α_1_ ALP box, with an α_1_ TMD rotation of 1.8°, suggesting that the two ALP pockets in the pentamer have a different molecular pharmacology. Global TMD alignment, on the other hand, highlights a greater tilt of the M2 helices with respect to the pore axis, collectively yielding an enlarged and more symmetric ion pore in our ALP-bound structure compared to that without ALP (**Figure 2e; Extended Data Figure 8)**. In particular, the sidechains of the 9’-Leu residues, which are crucial for channel gating, are rotated out of the pore in the presence of ALP (**Figure 2e**).

Neurosteroids achieve GABA_A_R potentiation by enhancing the ability of agonists to gate the channel^12, 37, 38^. Such enhancement must be due to allosteric rearrangements in the GABA-binding ECDs, which we indeed observe in our structure. Specifically, global TMD alignment reveals a concerted ∼2° (between 1.5° and 2.5°) counter-clockwise rotation of individual ECDs compared to the ALP-free structure when viewed from the extracellular side (**Extended Data Figure 8**). This likely accommodates expansion of the TMDs via interactions between the ECD Cys loops and TMD TM2-TM3 loops. Although GABA binding remains largely unchanged (**Extended Data Figure 8**), concerted ECD rotations may pose an additional energy barrier to GABA release, thus slowing its unbinding and increasing channel gating. Furthermore, because agonist-induced gating is known to be accompanied by counter-clockwise rotation of the ECDs^6^, our observed conformational changes are fully compatible with allosteric potentiation of n*α*_1_GABA_A_Rs by ALP. Thus, despite both molecular models in this structural comparison being in a desensitized state with the 2’ gate closed, ALP-induced remodeling of TMDs and ECDs explains how the receptor opens more readily in the presence of ALP. Furthermore, our data suggest that direct activation by neurosteroids is mediated via the same two binding pockets, as no additional ALP molecules were resolved in samples prepared with ALP concentrations as high as 5 μM.

Neurosteroids have unusually slow on- and off-rates compared to more hydrophilic ligands^39, 40^. This behavior has been attributed to their lipophilic nature and tendency to be enriched in the membrane^41^, but consideration of lipids in our ALP-bound structure offers an additional explanation for this phenomenon. An annulus of lipids with distorted acyl tails completely buries ALPs in their binding sites. In total, we resolve nine lipid-like molecules at the β_2_^+^/α_1_^-^ interface (^+^ denoting the principal face and – denoting the complementary face), three being less than 5 Å from ALP (**Figure 2f**). As a consequence, ALP molecules must coordinate with the motions of these annular lipids to secure an exit pathway from the pocket, and partially disassociated ALP molecules may effectively re-engage the receptor without leaving the pocket via the housing provided by these lipids.

In addition to their prevalence in the ALP binding pockets, lipids structurally engage the receptor at other sites. The greater TMD tilt in our ALP-bound structure creates five inter-subunit pockets near the center of the membrane’s plane. All five pockets, including two general anesthetic binding sites, are occupied with lipid tails bent like a snorkel (**Figure 2g**). Collectively, these lipids serve as small wedges that stabilize the expanded conformation of the TMD.

Both meta-one-Fab and ortho-one-Fab have ALP bound at their β_2_^+^/α_3_^*-^ or β_2_^*+^/α_3_^-^ pockets, demonstrating that neurosteroid binding at the β^+^/α^-^ interface is independent of subunit identity and arrangement within the pentamer (**Extended Data Figure 7**). Consistent with this notion, residues involved in binding ALP are conserved in all β and α subunits. Thus, we propose that neurosteroid potentiation of all n*α*_1_GABA_A_Rs with a β^+^/α^-^ interface involves a mechanism similar to the one we have described for ALP binding to the native tri-heteromeric α_1_β_2_γ_2_ receptor. Nevertheless, our structures suggest there may be differences in potency or efficacy at each of the neurosteroid binding sites. Although the sequences forming the immediate ALP pockets are identical, the W245 residue (α_1_ numbering) in other α subunits adopts a different sidechain conformer, and can serve as a longer and more effective lever for ALP to reshape the TMD and potentiate receptor activity, consistent with previous electrophysiology experiments^42, 43^.

Strikingly, we observed similar neurosteroid densities in the ZOL/GABA dataset in the absence of added neurosteroid, which are best modeled as ALP molecules (**Extended Data Figure 8**). Although we did not locate any distinct neurosteroid densities in the DID dataset, this is likely due to the inferior map resolution. Analysis of our purified ZOL/GABA sample by high performance liquid chromatography and mass spectrometry confirmed the absence of neurosteroid in the buffer and lipids used for protein purification. However, the same analysis uncovered 115 ng/mL (362 nM) of neurosteroid in the ZOL/GABA cryo-EM sample (containing ∼250 nM pentameric receptor), with more than 95% being ALP rather than another of the other three possible stereoisomers (**Extended Data Figure 9**). The nearly identical TMD configuration of the ALP/GABA and ZOL/GABA structures supports this chemical assignment (**Extended Date Figure 7**). Thus, endogenous ALP co-purifies with n*α*_1_GABA_A_Rs and its stoichiometric presence in our native receptor structures highlights its abundance in the brain and high affinity for n*α*_1_GABA_A_Rs relative to other endogenous neurosteroids.

### DID and ZOL augment GABA-induced rearrangements

GABA_A_Rs are the target of a range of insomnia medicines, including flurazepam and ZOL. To investigate the molecular effects of insomnia treatments on n*α*_1_GABA_A_Rs, we examined the interactions with either DID, the major metabolite of flurazepam^44^ (K_i_ 16.9 ± 1.7 nM), or ZOL (K_i_ 22.9 ± 2.7 nM) and n*α*_1_GABA_A_Rs (**Figure 3a and 3b**). Both compounds engage the receptor ECD at the α1^*+^/γ2^-^ interface, which is spatially equivalent to the GABA pockets, each sandwiched at a β^+^/α^-^ interface. The binding of DID in the two-Fab dataset is reminiscent of recombinant GABA_A_R structures in complex with diazepam/alprazolam^6^. DID makes extensive interactions with the receptor, including a hydrogen bond between its carbonyl and the α_1_-S204 sidechain; two hydrogen bonds between its A ring chloride and the α_1_-H101 and γ_2_-Ν60 sidechains; and several π-π/CH interactions with α_1_-F99, α_1_-Y159, α_1_-Y209, and γ_2_-Y58 (**Figure 3c**).

**Figure 3.**
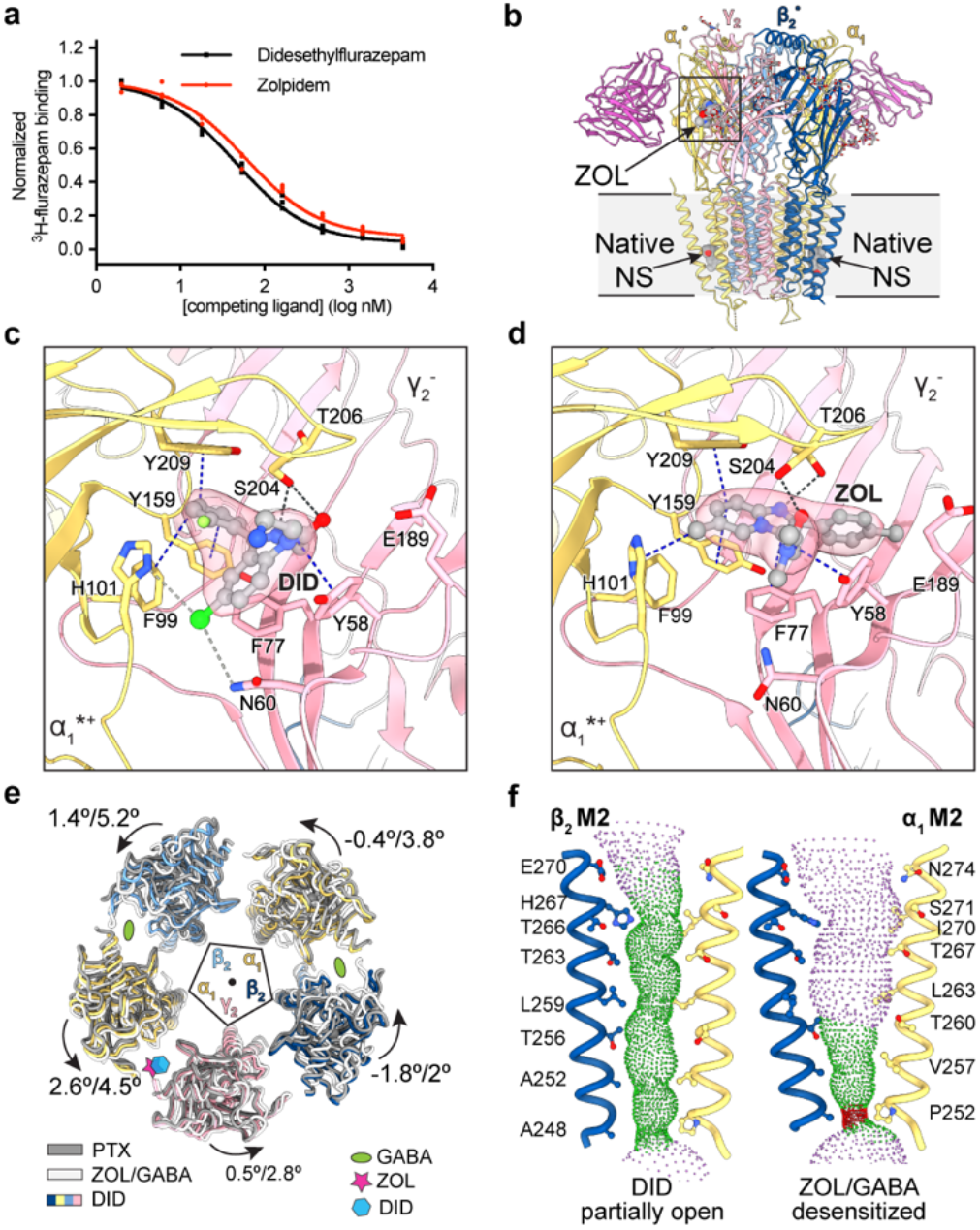
DID and ZOL binding propagates conformational changes to the TMD. **a**, Competitive radioligand binding assay for purified nα_1_GABA_A_Rs in complex with ZOL or DID. Individual data points are plotted along with the curve fitted with the one-site model. **b**, Structural overview of α_1_*-β_2_-α_1_-β_2_*-γ_2_ nα_1_GABA_A_R in complex with ZOL, GABA, and endogenous neurosteroid. **c, d**, Binding poses of DID and ZOL, and cryo-EM ligand density. Blue dashed lines, π-π/CH interactions; black dashed lines, hydrogen bonds less than 3.5 Å (acceptor to donor); gray dashed lines, weak hydrogen bonds between 3.5 Å and 4 Å. **e**, Impact of ligand binding on ECD arrangement. Structures superimposed based on global TMD. Individual ECDs displaced from center of pore by 15 Å for clarity. **f**, Pore profiles in DID/GABA structure and ZOL/GABA structure. Pore-delineating dots colored according to pore radius at that position: red, <1.8 Å; green, between 1.8 Å and 4 Å; blue, >4 Å.

ZOL binds to the α_1_^*+^/γ_2_^-^ ECD interface in tri-heteromeric α_1_β_2_γ_2_ receptors at roughly the same position as DID, but engages α_1_-H101 via π-CH interactions rather than a hydrogen bond. In addition, its amide oxygen forms a hydrogen bond with the α_1_-S204 sidechain, and the imidazole nitrogen forms a separate hydrogen bond with the α_1_-T206 sidechain (**Figure 3d**). We hypothesize that this hydrogen bond duet is preserved in interactions with α_2_ and α_3_ subunits but not the α_5_ subunit in which a threonine residue substitutes for S204. This difference would provide an explanation for the greater than ten-fold weaker affinity of ZOL for α_5_-containing receptors^45^. Like DID, ZOL forms π-π interactions with α_1_-Y159, α_1_-Y209, and γ_2_-Y58, as well as γ_2_-F77. The latter interaction explains why ZOL is more sensitive to the γ2-F77I mutation than diazepam^46^. During the preparation of this manuscript, a recombinant GABA_A_R structure in complex with ZOL was published^47^, revealing a similar binding pose for ZOL in the ECD. This structure also captured ZOL in the general anesthetic binding pockets at the β_2_^+^/α_1_^-^ TMD interface using a similar ZOL concentration to that used in our study. We hypothesize that remodeling of the TMDs by endogenous ALP prevented ZOL from binding to the general anesthetic pockets in n*α*_1_GABA_A_Rs.

Binding of DID or ZOL causes only moderate conformational changes in their binding pocket. We observed a slight opening of loop C due to a 1.2 Å displacement of the γ_2_-S205 Cα, as well as sidechain reorganization of α_1_-H101, γ_2_-Y58, γ_2_-Ν60, and γ_2_-F77, which enlarges the pocket to accommodate the ligand. These subtle changes suggest that ZOL-like medications (Z-drugs) potentiate GABA_A_Rs via a benzodiazepine-like mechanism, namely, strengthening of the α_1_^*+^/γ_2_^-^ interface and facilitation of GABA-induced ECD rotation^6^. Indeed, when the TMDs of the ZOL/GABA structure were aligned to the closed, resting structure^6^, the ECDs showed concerted counterclockwise rotations ranging from 2 to 5° for individual ECD centers of mass (**Figure 3e**).

We also observed ZOL binding to the α_1_^*+^/γ_2_^-^ (ortho-one-Fab) and α_3_^*+^/γ_2_ (meta-one-Fab) ECD interfaces. Despite sequence differences, the immediate α3^*+^/γ2 pocket shares the same chemical environment as the α_1_^*+^/γ_2_^-^ pocket, but the sidechains adopt different conformations. Accordingly, we observed significantly different structural consequences of ZOL binding in the α_3_^*+^/γ_2_ pocket, including a binding pose closer to loop C on the α_3_ subunit and concerted shifts of the ligand and the protein (**Extended Data Figure 7**). As mentioned above, our data suggest that α_2/3/5_ subunits have a greater intrinsic bend between their ECD and TMD than α_1_ subunits, causing different global rearrangements when incorporated into the pentamer. Although this variation in ECD/TMD coupling causes relatively small structural perturbations to orthosteric and allosteric ligand binding, it has the potential to affect channel gating and ligand modulation, which depend on inter-domain cross-talk.

Intriguingly, the minimum pore radius in the DID structure is 2 Å, large enough to pass dehydrated Cl^-^ with a radius of 1.81 Å^48^ (**Figure 3f**). The capture of this potentially conductive state, which has not been observed before, is likely due to prevention of GABA-induced desensitization (GABA was omitted during DID sample preparation) and potentiation by DID (micromolar concentrations of benzodiazepines potentiate GABA_A_Rs^49, 50^). Accordingly, we observed incomplete loop C closure – the structural hallmark of GABA-dependent allostery – in the GABA binding pockets in the DID structure.

## Conclusion

Our study reveals that n*α*_1_GABA_A_Rs comprise three structural populations: tri-heteromeric α_1_β_2_γ_2_ receptors that constitute half the total population and two distinct assemblies containing one α_1_ subunit and one α_2/3/5_ subunit. Because only three distinct receptor assemblies were identified from a total of 72 possible arrangements of two α, one β, and one γ subunit^18^, we propose that neuronal assembly of n*α*_1_GABA_A_Rs is a highly regulated process. The finding of exclusively α-β-α-β-γ_2_ assemblies is in contrast to the diversity of subunit combinations that are formed in recombinant expression systems, where β-β interfaces and assemblies with two γ subunits are observed^18^. Although our cryo-EM study validates the use of tri-heteromeric α_1_β_2_γ_2_ receptors as a model of n*α*_1_GABA_A_R pharmacology, it also reveals the prevalence of mixed α subunit receptors in the brain and challenges the conventional practice of classifying GABA_A_Rs according to α subunits.

The molecular structures of ligand-n*α*_1_GABA_A_R complexes have revealed the binding poses of the postpartum depression medication, allopregnanolone, and two insomnia drugs, didesethylflurazepam and zolpidem. Our work also highlights the conformational changes induced by neurosteroid binding to native receptors and thus the structural basis for neurosteroid-dependent positive modulation (**Figure 4**). Finally, the serendipitous finding that endogenous neurosteroids remain bound to n*α*_1_GABA_A_Rs after isolation and purification emphasizes the importance of considering background neurosteroid modulation when investigating the pharmacology of n*α*_1_GABA_A_Rs.

**Figure 4.**
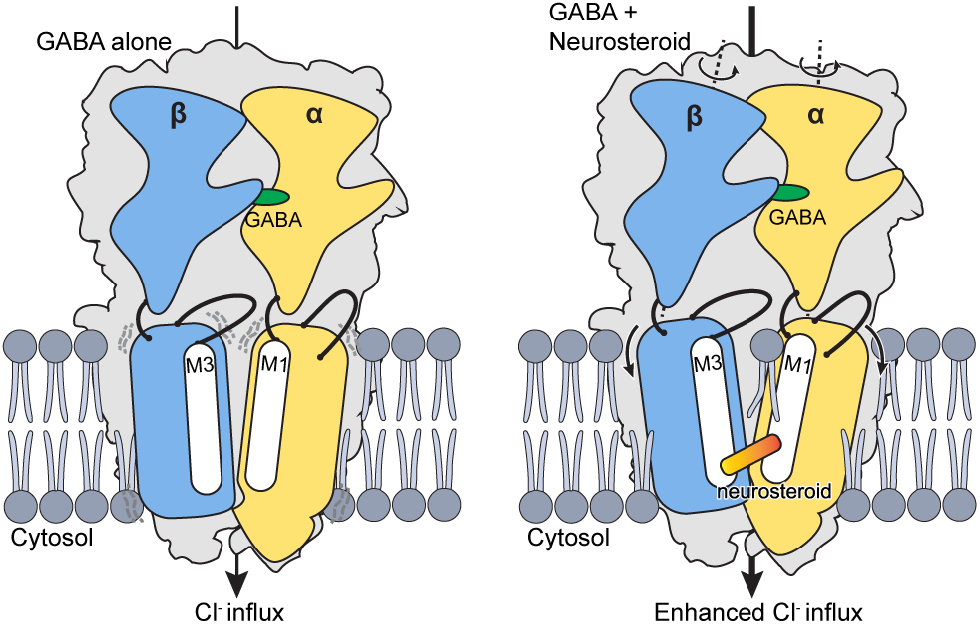
Mechanism of neurosteroid potentiation. Binding of GABA to nα_1_GABA_A_Rs induces rotation of the ECD and opening of TMD pore, allowing Cl^-^ ion influx. Neurosteroid bound in the β^+^/α^-^ TMD pocket serves as a diagonal brace to stabilize the open conformation of the TMD and induce additional counter-clockwise rotation of individual ECDs. Annular lipid molecules position their acyl tails between the M3 and M1 helices of adjacent subunits, contributing further stabilization. The net effect of neurosteroid binding is enhanced Cl^-^ influx.

## Acknowledgements

We thank J. Luo and A. DeBarber for mass-spec neurosteroid analysis, J. Guidry for mass-spec protein identification, D. Claxton and D. Cawley for the monoclonal antibody. We thank the use of OHSU Bioanalytical Shared Resource/Pharmacokinetics Core Facility expertise and instrumentation (Research Resource Identifier (RRID): SCR_009963). We thank OHSU Multiscale Microscopy Core (MMC), the Pacific Northwest Cryo-EM Center (PNCC), and the cryo-EM facility at Janelia research campus for microscope use. C.S. thanks J. Myers (PNCC) and C. López (MMC) for cryo-EM training. PNCC is supported by NIH grant U24GM129547 and accessed through EMSL (grid.436923.9), a DOE Office of Science User Facility sponsored by the Office of Biological and Environmental Research. This work was supported by NIH grant 5R01GM10040 to E.G. and E.G. is an investigator of the Howard Hughes Medical Institute.

## Author Information

Hongtao Zhu

Present address: Institute of Physics, Chinese Academy of Sciences, Beijing, China

## Authors and Affiliations

**Vollum Institute, Oregon Health and Science University, Portland, OR, USA**

Chang Sun, Hongtao Zhu, Sarah Clark, Eric Gouaux

## Contributions

C.S. and E.G. designed the project. C.S. prepared cryo-EM samples, carried out biochemical characterizations. C.S. and S.C. performed single-molecule photobleaching experiments. C.S. and H.Z. carried out the cryo-EM data analysis and C.S. built the molecular models. C.S. and E.G. wrote the manuscript.

## Data availability

The cryo-EM maps and coordinates for the native GABA receptor in complex with didesethylflurazepam and endogenous GABA (two-Fab-DID) have been deposited in the Electron Microscopy Data Bank (EMDB) under accession number EMD-29728 and in the Protein Data Bank (PDB) under accession code 8G4O. The cryo-EM maps and coordinates for the native GABA receptor in complex with zolpidem, GABA, and endogenous neurosteroids have been deposited and accessed via EMD-39727/8G4N (two-Fab-ZOL), EMD-29743/8G5H (ortho-one-Fab-ZOL), EMD-29742/8G5G (meta-one-Fab-ZOL). The cryo-EM maps and coordinates for the native GABA receptor in complex with GABA, and allopregnanolone have been deposited and accessed via EMD-29350/8FOI (two-Fab-ALP), EMD-29741/8G5F (ortho-one-Fab-ALP), EMD-29733/8G4X (meta-one-Fab-ALP).

## Methods

### Expression and purification of the α_1_ specific 8E3-GFP Fab

The α1-specific mouse monoclonal antibody 8E3 was generated and screened as previously described^1^. The coding sequences of 8E3 Fab light and heavy chains were determined from hybridoma mRNA, and a construct to express the Fab portion of the antibody was designed by including sequences to encode an N-terminal GP64 signal peptide. Codons were optimized for expression in insect cells. To facilitate recombinant antibody detection and purification, a 3C-cleavage sequence, an EGFP gene, and a twin-strep II tag were added to the C-terminus of the heavy chain. Synthetic genes for both chains were then cloned into the pFastBac-Dual vector under the polyhedrin promoter. The recombinant baculovirus was prepared as previously described^2^. Sf9 cells at a density of 3 million per mL were infected with the recombinant baculovirus, with a multiplicity of infection of 2, and further cultured for 96 hours at 20° C. The antibody-containing supernatant was collected by a 20-minute centrifugation at 5,000 g and then the pH was adjusted to 8 with 30 mM Tris base, incubated in the cold room overnight to allow non-Fab protein precipitation, and clarified by another 20-minute centrifugation at 5,000 g. The supernatant was concentrated and buffer-exchanged three times with TBS (20 mM Tris 150 mM NaCl pH 8) using a tangential-flow concentrator equipped with a 15-kDa filter. The concentrated supernatant was then loaded onto a 15-mL streptactin column, which was washed with at least 20 column volumes of TBS and eluted with 5 mM desthiobiotin in TBS. Selected fractions were pooled, concentrated, and buffer exchanged to TBS using microconcentrators with a 50-kDa cutoff. Concentrated 8E3-GFP Fab (∼100 μM) was aliquoted and stored at –80 ° C until use.

### Purification of nα_1_GABA_A_Rs from mouse brains

One-month-old BL/6 mice of mixed sex (∼50 mice per preparation) were used for native receptor isolation. The mice were first euthanized and decapitated. The whole brain was isolated from the skull using a laboratory micro spatula and stored in ice-cold TBS. Cerebella were removed from the whole brain, frozen in liquid nitrogen, and stored at -80 ° C for a separate study. After being washed twice with ice-cold TBS, brain tissue was resuspended with ice-cold TBS (1 mL per brain) supplemented with 0.2 mM phenylmethyl sulfonyl fluoride (PMSF). The suspension was processed with a loose-fit Potter-Elvehjem homogenizer for 20 full up-and-down strokes and further sonicated (1 min per 50 mL) at a setting of 6, typically at a 40 W output. The suspension was centrifuged at 10,000 g for 10 minutes, resulting in a hard pellet of mainly the nuclear fraction and a “runny” soft pellet containing a significant amount of nα_1_GABA_A_Rs. The supernatant was further centrifuged at 200,000 g for 45 minutes to pellet the membranes. About 0.1 g of hard pellet and 0.2 g of soft pellet were obtained from one mouse brain. These membrane pellets were resuspended with an equal volume of TBS buffer containing protease inhibitors (aprotinin/leupeptin/pepstatin A/PMSF). If not used right away, the 50% membrane suspension was supplemented with 10% glycerol and snap frozen in liquid nitrogen.

The following membrane solubilization and affinity chromatography were all carried out at 4 ° C. First, MNG/CHS (10:1 w/w) stock (10% w/v) was diluted in TBS buffer containing protease inhibitors to 2.5%. Then, one volume of the 50% membrane suspension was mixed with two volumes of the diluted detergent stock and incubated for 1 hour on a platform rocker, which routinely resulted in the solubilization of ∼60% of the α_1_ subunit present in the tissue, estimated based on Western blot (**Extended Data Figure 1**). Next, Biolock solution was added at 0.1 mL per brain to quench the naturally biotinylated proteins, and the mixture was clarified by centrifugation at 200,000 g for 1 hour. Finally, the 8E3-GFP Fab was added to the solubilized membrane to a concentration between 60 nM and 100 nM. After 1 hour incubation, 3 mL of pre-equilibrated streptactin resin was added to bind the 8E3-GFP Fab and associated nα_1_GABA_A_Rs, for 2 hours in batch mode.

### On-column nanodisc reconstitution

MSP2N2^3^ or a recently engineered MSP1E3D1 variant, CSE3^4^, was used for on-column MSP nanodisc reconstitution. The affinity resin, bound with receptor complexes, was washed in batches, first with 20-CV of ice-cold TBS, then with 20-CV of TBS containing 0.05% MNG and 0.01% brain polar lipid (Avanti). During this wash, 40 nmole MSP2N2 and 3.2 μmole POPC:bovine brain extract (Sigma) (85:15) lipids, or 40 nmole CSE3 and 4.8 μmole lipids were mixed to a final volume of 1 mL in TBS and incubated at room temperature for 30 minutes. The beads were transferred to an empty Econo-Pac gravity flow column to drain the buffer. Then the 1 mL pre-incubated MSP:lipids were added and incubated for 1.5 hours. Next, biobeads were added to a 20x weight excess to the MNG detergent. The mixture was incubated with a rotator in the cold room for at least 4 hours. The biobead/resin mixture was washed with 20-CV of ice-cold buffer to remove unbound empty nanodiscs.

Two approaches were used to elute reconstituted nanodiscs: competitive ligand elution and protease cleavage. For ligand elution, 0.5 CV 5 mM desthiobiotin dissolved in TBS was incubated with the streptactin superflow resin for 10 minutes before gravity elution, which was repeated for a total of 6 times. In the case of 3C cleavage, 0.1 mg 3C protease was first diluted to 50 μg/mL with 2 mL TBS and added to the resin. After a 2-hour incubation in the cold room, the elution was collected, and the column was further washed three times with 2 mL TBS to improve the protein yield. 3C protease cleavage offered better protein purity and was used for the zolpidem and the allopregnanolone samples. Pooled elution was concentrated to about 0.5 mL using a 50-kDa cutoff centricon, regardless of the elution methods. The concentrated sample was then injected into a Superose 6 column pre-equilibrated with TBS supplemented with 1 mM GABA and other ligands. Selected fractions corresponding to the nα_1_GABA_A_R:Fab complex were combined and concentrated to about 0.1 mg/mL using a centricon with a 50-kDa cutoff.

### Mass-spec protein identification

The protein mass-spec analysis was carried out as previously described.^5^ The native receptor samples were diluted into 100 μl 1% SDS, reduced with Tris (2-carboxyethyl) phosphine hydrochloride (TCEP), and alkylated with iodoacetamide. Proteins were then extracted with methanol-chloroform, mixed with 30 μl of 20 μg/mL trypsin dissolved in 50 mM ammonium bicarbonate, and incubated overnight at 37 °C. The next day, the solvent from the trypsin digestion was evaporated using a speed vac. The resulting pellet was resuspended in 20 μl of 2% acetonitrile (ACN) and 0.1% formic acid (FA) for LC-MS. The sample was run on a Dionex U3000 nanoflow system coupled to a Thermo Fusion mass spectrometer. Each sample was subjected to a 65-min chromatographic method using a gradient from 2–25% acetonitrile in 0.1% formic acid (ACN/FA) for 16 min; from 25% to 35% ACN/FA for an additional 15 min, from 35% to 50% ACN/FA for an additional 4 min, a step to 90% ACN/FA for 4 min and a re-equilibration into 2% ACN/FA. Chromatography was carried out in a ‘trap-and-load’ format using a PicoChip source (New Objective); trap column C18 PepMap 100, 5 µm, 100 A, and the separation column was PicoChip REPROSIL-Pur C18-AQ, 3 µm, 120 A, 105 mm. The entire run was at a flow rate of 0.3 µl/min. Electrospray was achieved at 1.9 kV. The MS1 scans were performed in the Orbitrap with a resolution of 240,000. Data-dependent MS2 scans were performed in the Orbitrap using High Energy Collision Dissociation (HCD) of 30% using a resolution of 30,000. Data analysis was performed using Proteome Discoverer 2.3 using SEQUEST HT scoring. The static modification included dynamic modification of methionine oxidation (+15.9949) and a fixed modification of cysteines alkylation (+57.021). Parent ion tolerance was 10 ppm, fragment mass tolerance was 0.02 Da, and the maximum number of missed cleavages was set to 2. Only high-scoring peptides were considered, using a false discovery rate (FDR) of 1%.

### Isotope-dilution quantification of allopregnanolone using LC-MS/MS

Neurosteroids (allopregnanolone, epipregnanolone, isopregnanolone, pregnanolone) and isotope-labeled internal standard allopregnanolone-d5 were purchased from Toronto Research Chemicals (Toronto, ON, Canada). The O-(3-trimethylammonium-propyl) hydroxylamine quaternary amonoxy (QAO) reagent used for derivatization was in the form of Amplifex Keto reagent kit from AB Sciex (Framingham, MA). Solvents for liquid chromatography-tandem mass spectrometry (LC-MS/MS) analysis were from VWR (Tualatin, OR).

Neurosteroid stocks and internal standard (INST) were prepared in methanol. Stocks (5 μL) and the INST allopregnanolone-d5 (5 μL) were mixed with PBS (95 μL) to prepare standard samples with final concentrations ranging from 0.05 to 100 ng/ml. All standards and samples were treated with 1000 µl of acetonitrile, vortexed and mixed using Benchmark Multi-Thermo heat/shaker at 1500 rpm at 22°C for 5 mins, and centrifuged to remove protein at 12,000 g for 5 mins. The supernatant was dried under vacuum and then treated with 75 µl of derivatization reagent. The keto moiety was derivatized with QAO reagent to form a cationic oxime derivative to enable highly sensitive LC-ESI-MS/MS quantification of neurosteroids. The working derivatization reagent was prepared according to vendor instructions. The derivatized samples were diluted 1:4 with 5% acetic acid in methanol before LC-MS/MS analysis. The supernatant was placed in sample vials for analysis by LC-MS/MS using an injection volume of 5 µl. The lower limit of quantification of allopregnanolone was 75 pg/ml, with an accuracy of 101% and a precision of 2.2%.

The samples with INST were analyzed using a Sciex 4000 QTRAP hybrid/triple quadrupole linear ion trap mass spectrometer (Foster City, CA) with electrospray ionization (ESI) in the positive mode. The mass spectrometer was interfaced to a Shimadzu HPLC system (Columbia, MD) with SIL-20AC XR auto-sampler, LC-20AD XR LC pumps, and CTO-20AC column oven. Compounds were quantified with multiple reaction monitoring (MRM). The MS/MS transitions used were optimized by infusion of pure derivatized compounds with method settings, as presented in the table below. The bold transitions were used for quantification, with other transitions used for peak qualification to ensure method specificity. Allopregnanolone was separated from interferents using a Luna 5u C8(2) 50×2 mm column (Phenomenex) kept at 35 °C using a column oven. The gradient mobile phase was delivered at a flow rate of 0.8 ml/min and consisted of two solvents: solvent A (0.1% formic acid in water) and solvent B (0.1% formic acid in acetonitrile). The initial concentration of solvent B was 20%, followed by a linear increase to 60% B in 10 min, then to 95% B in 0.1 min, held for 3 minutes, decreased back to starting 20% B over 0.1 min, and then held for 2 min. The retention time was 3.99 min for allopregnanolone and pregnanolone, 3.64 min for isopregnanolone, and 3.61 min for epipregnanolone. Data were acquired using Analyst 1.6.2 and analyzed with MultiQuant 3.0.3 software.

To further distinguish allopregnanolone and pregnanolone, a different HPLC condition was used. In this case, a Poroshell 120 EC-C18 100×2.1 mm 2.7um column (Agilent) was kept at 35 °C using a column oven. The gradient mobile phase was delivered at a flow rate of 0.4 ml/min (0–5.9min), 0.2ml/min (6.0–8.9min), and 0.4ml/min (9–15min), and consisted of two solvents, A: 0.1% formic acid in water, B: 0.1% formic acid in acetonitrile. The initial concentration of solvent B was 30%, followed by a linear increase to 52% B in 6.5 min, held for 2.5min, then to 95% B in 0.1 min, held for 2.9 minutes, decreased back to starting 30% B over 0.1 min, and then held for 2.9 min. The retention time for allopregnanolone was 6.4 min, pregnenolone was 6.2 min, and 3*α*-allopregnanolone-d5 was 6.3 min.

| Q1 mass | Q3 mass | Dwell Time (msec) | ID | DP | EP | CE | CXP |
| --- | --- | --- | --- | --- | --- | --- | --- |
| 433.3 | 374.3 | 200 | QAO-ALLOPREGNANOLONE | 71 | 10 | 35 | 14 |
| 433.3 | 58 | 200 | QAO- ALLOPREGNANOLONE | 71 | 10 | 129 | 8 |
| **433.3** | **126.1** | **200** | **QAO- ALLOPREGNANOLONE** | **71** | **10** | **53** | **12** |
| **438.3** | **126** | **200** | **QAO- ALLOPREGNANOLONE-d5** | **76** | **10** | **55** | **18** |

### Single-molecule photobleaching of nα_1_GABA_A_R-Fab complexes

Coverslips and glass slides were extensively cleaned, passivated, and coated with methoxy polyethylene glycol (mPEG) and 2% biotinylated PEG as previously described^6^. A flow chamber was created by drilling 0.75 mm holes in the quartz slide and placing double-sided tape between the holes. A coverslip was placed on top of the slide, and the edges were sealed with epoxy, creating small flow chambers. A concentration of 0.25 mg/mL streptavidin was then applied to the slide, incubated for 5 minutes, and washed off with buffer consisting of 50 mM Tris, 50 mM NaCl and 0.25 mg/mL bovine serum albumin (BSA), pH 8.0. Biotinylated anti-GFP nanobody at 7.5 µg/mL was applied to the slide, incubated for 10 minutes, and washed off with 30 µL buffer A (20 mM Tris, 150 mM NaCl, pH 8) supplemented with 0.2 mg/mL BSA. nα_1_GABA_A_R-Fab complexes in nanodiscs were eluted from the streptactin-XT resin with biotin instead of 3C protease cleavage to preserve the GFP moiety. The sample was further FSEC-purified, and the peak corresponding to the complex was hand collected, which separated the native receptor from free Fab. The sample was diluted 1:30 to about 50 pM based on fluorescence quantitation, applied to the chamber, and incubated for 5 minutes before being washed off with 30 µL of buffer A. The chamber was immediately imaged using a Leica DMi8 TIRF microscope with an oil-immersion 100x objective. Images were captured using a back-illuminated EMCCD camera (Andor iXon Ultra 888) with a 133 x 133 µm imaging area and a 13 µm pixel size. This 13 µm pixel size corresponds to 130 nm on the sample due to the 100x objective.

Photobleaching movies were acquired by exposing the imaging area for 180 seconds. Single-molecule fluorescence time traces of nα_1_GABA_A_R-Fab were generated using a custom python script. Each trace was manually scored as having one to three bleaching steps or was discarded if no clean bleaching steps could be identified. A total of ∼450 molecules were evaluated from three separate movies. Scoring was verified by assessing the intensity of the spot; on average, the molecules that bleached in 2 steps were twice as bright as those that bleached in 1 step.

### Scintillation proximation assay

YSI Copper SPA beads from PerkinElmer were used to capture the nα_1_GABA_A_R in nanodisc via the MSP His-tag. Tritiated flunitrazepam from PerkinElmer was used as the radioligand, and clorazepate was used as the competing ligand to estimate background. During the ligand binding assay setup, nα_1_GABA_A_R in nanodisc was first mixed with SPA beads and radioligand (2x bead) while the ligand of different concentrations (2x ligand) and competing ligand (2x background) were prepared using serial dilution. Then, an equal volume of 2x bead was mixed with 2x ligand (in triplicate) or 2x background in a 96-well plate. The final concentrations were 0.5 mg/mL for SPA beads, ∼1 nM for native receptors, 10 nM for ^3^H-flunitrazepam, and 0.5 mM for clorazepate in the background wells only. The plate was then read with a MicroBeta TriLux after a 2-hour incubation. Specific counts were then imported into GraphPad and analyzed using a one-site competition model.

### Negative-stain electron microscopy

Purified nα_1_GABA_A_R:Fab complex in nanodiscs was first diluted with TBS to a concentration of ∼0.05 mg/mL. Continuous carbon grids were glow-discharged for 60 seconds at a current of 15 mA. A protein sample (5 μL) was applied to the carbon side of the grid held with a fine-tip tweezer and incubated for 10–30 seconds. The excessive sample was then wicked away from the side with a small piece of filter paper. The grid was quickly washed with 5 μL deionized water, followed by side-wicking, which was repeated for a total of three times. Immediately afterward, the grid was incubated with 5 μL 0.75% uranium formate for 45 seconds, wicked several times from the side, and dried for at least 2 minutes at room temperature.

### Cryo-EM sample preparation and data acquisition

We employed a specific setup to prepare grids under different buffer and ligand conditions. First, buffers containing 10x ligand or additive were first prepared and dispensed in 0.5 μL aliquot into PCR tubes. Then, 5 μL purified nα_1_GABA_A_R:Fab complex was added and quickly mixed by pipetting. Within 10 seconds, a 2.5 μL sample was applied to a glow-discharged (30 seconds at 15 mA) 200 mesh gold Quantifoil 2/1 grid overlaid with 2-nm continuous carbon and incubated for 30 seconds. The grid was blotted with a Mark IV Vitrobot under 100% humidity at 16 °C and flash-frozen in liquid ethane. For the didesethylflurazepam sample, no GABA was included during the purification, and the didesethylflurazepam (2 μM) was added prior to vitrification. For the zolpidem sample, 1 mM GABA was included throughout the purification, and 5 μM zolpidem was added prior to vitrification using the above-mentioned PCR tube method. For the allopregnanolone sample, 1 mM GABA and 5 μM allopregnanolone were included from the membrane solubilization to the final size-exclusion chromatography.

Cryo-EM data were collected on a 300-keV Titan Krios equipped with a BioQuantum energy filter at either PNCC or the Janelia cryo-EM facility. Data acquisition was automated using serialEM: defoci ranged between 0.9 to 2.5 μm, holes with suitable ice thickness were selected with the hole finder and combined to produce multishot-multihole targets, which allowed the acquisition of six movies per hole in each of the neighboring nine holes. These movies were captured with a K3 direct electron detector. A total dose of 50 electron/Å^2^ was fractionated into 40 frames, with a dose rate of about 15 electron/(pixel*second) for non-CDS mode or 7 electron/(pixel*second) for CDS mode (**Extended Data Table 1**).

### Cryo-EM data analysis

Super-resolution movies were imported to cryosparc^7^ v 3.3.1 and motion corrected using cryosparc’s patch motion correction with the output Fourier cropping factor set to ½. Initial contrast transfer function (CTF) parameters were then calculated using cryosparc’s patch CTF estimation. For each dataset, 2D class averages of particles picked by glob-picker from ∼1000 micrographs were used as templates for the template picker. One round of 2D classification and several rounds of heterogeneous refinement seeded with *ab initio* models generated within cryosparc were used to select GABA_A_R particles, ranging from 4 to 6 million particles for our datasets. A non-uniform refinement (NU-Refinement) was performed to align these particles to a consensus structure. Two downstream strategies were used for our datasets, as subsequently described.

#### Data processing strategy #1

Bin 1 GABA_A_R particles, both images (360×360) and the star file converted using pyem^8^, were ported into RELION^9^ 3.1. Then a 3D auto refinement job with local search (angular sampling of 1.8 degree) was carried out to fine tune the particle poses in RELION. The refined structure, similar to that generated by cryosparc, had relatively weaker γ subunit transmembrane helices, which was reported previously^10^. To tackle this issue, we prepared a nanodisc mask in Chimera^11^ and carried out 3D classification without alignment (15 classes, T=20) using that mask. The 3D classification can robustly give classes with much stronger transmembrane helices of the γ subunit. Those selected particles were imported into cryosparc and further refined using NU-Refinement with both defocus refinement and per-group CTF refinement options turned on. The consensus structure was a two-Fab bound structure, but earlier data processing revealed one-Fab species’ presence. Therefore, a 3D classification job was used with a mask focusing on the two binding sites of 8E3 Fab to isolate the one-Fab species. The one-Fab and two-Fab particles were separately refined with NU-Refinement and further refined with local refinement.

#### Data processing strategy #2

In this strategy, the heterogeneity in Fab binding was addressed upstream in the data processing pipeline. Like strategy #1, GABA_A_R particles, at bin3 or 120×120, were imported into RELION for focused 3D classification. A reverse mask was prepared in cryosparc which only excluded the transmembrane domain to allow for Fab binding at all possible positions. 3D classification (10 classes, T=20) gave clear two-Fab and one-Fab classes, and classes with incomplete Fab. Further 3D classification on these incomplete Fab particles produced only incomplete Fab classes, which led us to believe they were damaged particles and should be excluded from downstream processing. The two-Fab particles and the one-Fab particles, on the other hand, were imported into cryosparc, re-extracted at bin1, and separately refined with NU-Refinement. Still, we saw weak transmembrane helices for the γ subunit for the one-Fab and the two-Fab populations. To tackle this issue, instead of the 3D classification in RELION, we used the 3D classification (beta) job in cryosparc with a nanodisc mask, which was less robust but faster. Classes with stronger transmembrane helices were then combined and refined with NU-Refinement and finished with local refinement.

Global sharpening worked sub-optimally for our nα_1_GABA_A_R structures because of the local resolution variation and the lower signal-to-noise ratio for the transmembrane domain. The best method to sharpen our maps was achieved with LocScale^12^, which was used to represent some of our structures in **Figure 1**. DeepEMhancer^13^ can yield comparable sharpening for the protein but not for the annulus lipids.

### Subunit identification, model building, refinement, and validation

Due to the subunit specificity of 8E3-Fab, the subunit with 8E3-Fab bound is defined as α_1_. The remaining subunits can be easily classified as α, β, or γ from each subunit’s characteristic *N*-linked glycosylation patterns. It was clear that all 3D classes obtained are α-β-α-β-γ, clockwise, when viewed from the extracellular side of the membrane. Given the relative subunit abundance from earlier studies, we used α_1_-β_2_-α_1_-β_2_-γ_2_ as the starting model of the two-Fab class. We then examined the cryo-EM density maps to test our assignment in the context of sequence information. Specifically, we looked at regions where the sidechain can be unambiguously assigned and positions where a difference of more than 3 carbon atoms or one sulfur atom was found within the subunit group. Regarding the non-α_1_ α subunit in the one-Fab classes, we further limited our scrutiny to positions showing no significant conformational differences in the corresponding two-Fab structure to ensure the observed density difference was caused by the chemical identity of underlying residues.

For each dataset, the two-Fab bound nα_1_GABA_A_R model was built first. The starting structures used were Alpha-fold^14^ models of mouse GABA_A_R subunits and the best 8E3 Fab model generated with Rosetta^15^. These individual chains were first docked into the unsharpened cryo-EM density maps using chimera’s fit-in-map tool to assemble the full receptor-Fab complex. The full complex was then edited to remove unresolved portions and refined extensively to achieve better model-map agreement in Coot^16^. *N*-glycosylation was modeled using Coot’s carbohydrate module. Lipid and lipid-like molecules, including POPC, PIP2, dodecane, and octane, were modeled using the CCP4 monomer library. New ligands included in this study, including their optimized geometry and constraint, were generated using phenix.elbow^17^. After the initial modeling, multiple runs of phenix.real_space_refinement^18^ and editing in Coot were carried out to improve the model quality.

The optimized two-Fab GABA_A_R structure was used as the starting model for one-Fab GABA_A_R structures. Although the one-Fab population likely consists of a mix of α_2/3/5_ subunits, we decided to use α_3_ for the modeling because of its best overall agreement with the density maps. The two-Fab structure was first docked into the one-Fab cryo-EM map using the “fit in map” tool of chimerax^19^. Then the aligned structure was edited in Coot to remove the extra-Fab, replaced and renumbered the α_1_ sequence with the α3 sequence. This edited structure was further fitted and refined in Coot, first with secondary structure restraints generated with ProSMART^20^, and then without the restraints. Furthermore, certain residues and lipids were removed due to less clear density, and the glycosylation trees were remodeled. Similarly, this initial model was subjected to multiple runs of phenix.real_space_refinement and editing in Coot.

### Animal use statement

Mouse carcass donated from other labs of the Vollum Institute were used to establish and optimize the native GABA_A_ receptor isolation workflow. The quantity of purified native receptor from each mouse was estimated using the fluorescence from the recombinant antibody fragment, which was then extrapolated to give the minimum number required for cryo-EM and biochemical analysis. For each native GABA_A_ receptor preparation, 50 one-month-old (4–6 weeks) C57BL/6 mice (both male and female) were ordered from Charles River Laboratories. No randomization, blinding or experimental manipulations were performed on these animals. All mice were euthanized under Institutional Animal Care and Use Committee (IACUC) protocols, consistent with the recommendations of the Panel on Euthanasia of the American Veterinary Medical Association (AVMA) and carried out only by members of the E.G. laboratory approved on IACUC protocol TR03_IP00000905.

### Cell line statement

Sf9 cells for generation of baculovirus and expression of recombinant antibody fragment are from Thermo Fisher (12659017, lot 421973). The cells were not authenticated experimentally for these studies. The cells were tested negative for Mycoplasma contamination using the CELLshipper Mycoplasma Detection Kit M-100 from Bionique.

## Extended Data Figures

**Extended Data Figure 1.**
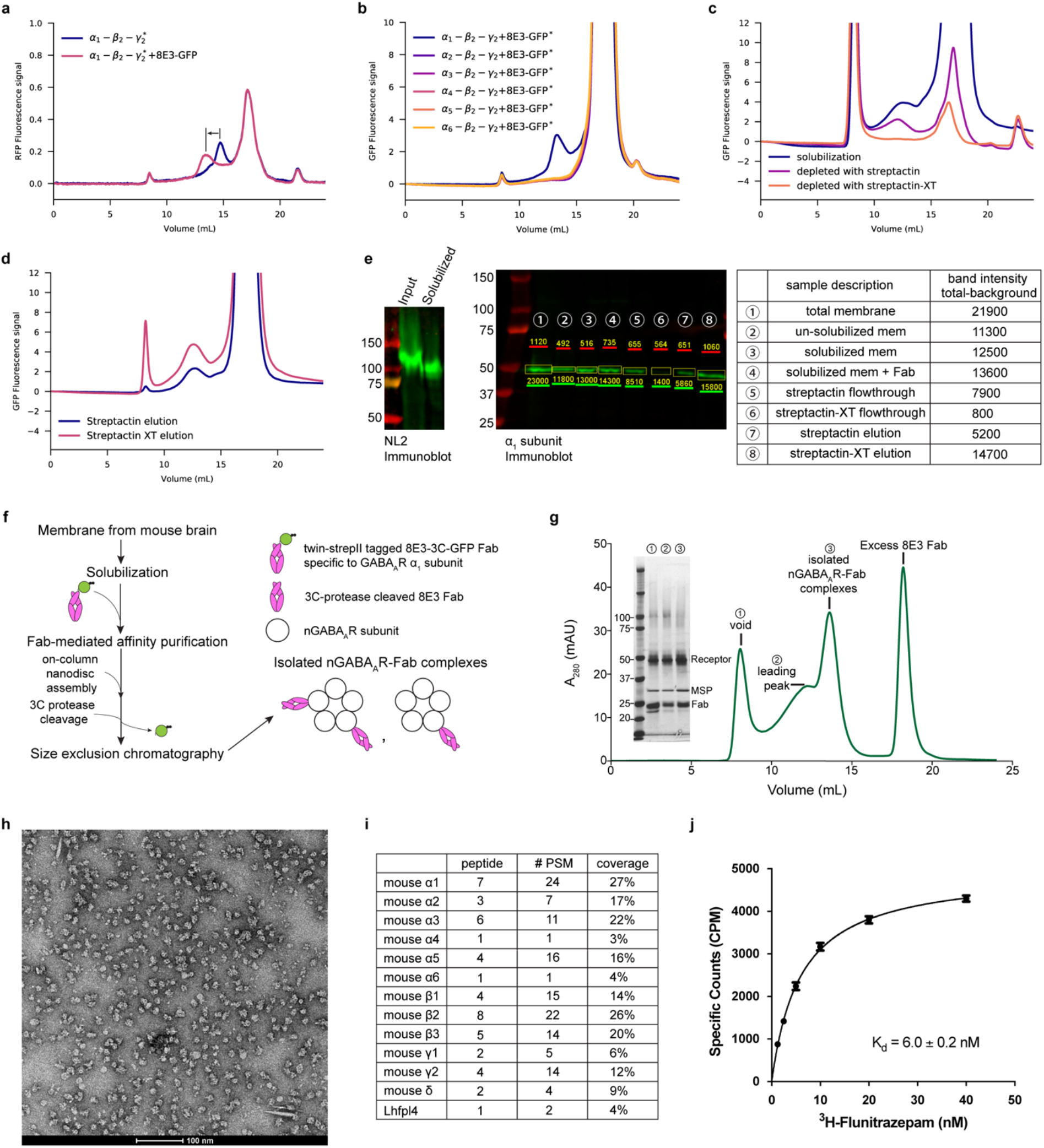
Biochemical characterization of native receptor isolation from mouse brain using an engineered Fab fragment. **a, b**, expression of tri-heteromeric GABA_A_Rs with different α subunits and binding test with the engineered 8E3-GFP Fab monitored with fluorescence-detection size-exclusion chromatography (FSEC). The signal is from the fusion-red protein inserted into the intracellular loop of the γ2 subunit in panel a or the GFP of the 8E3-GFP Fab in panel b. c, d, FSEC traces demonstrating the superior capturing efficiency and protein yield of streptactin-XT resin. e, Western blot analysis of steps during the native receptor purification. The neuroligin 2 (NL2) immunoblot shows robust solubilization of the inhibitory synapse marker NL2. The α_1_ subunit immunoblot shows quantitation of the α1 subunit during membrane solubilization, affinity capturing, and elution steps. f, the workflow of nα_1_GABA_A_Rs purification from mouse brain. g, size-exclusion chromatography (SEC) of nα_1_GABA_A_RS and silver-stain SDS-PAGE analysis of different SEC fractions. h, negative-staining electron microscopy images of protein samples from the pentameric peak. i, identified proteins related to GABA receptors from the pentameric peak using mass spectrometry. j, scintillation proximity assay of the pentameric peak fraction with ^3^H-Flunitrazepam.

**Extended Data Figure 2.**
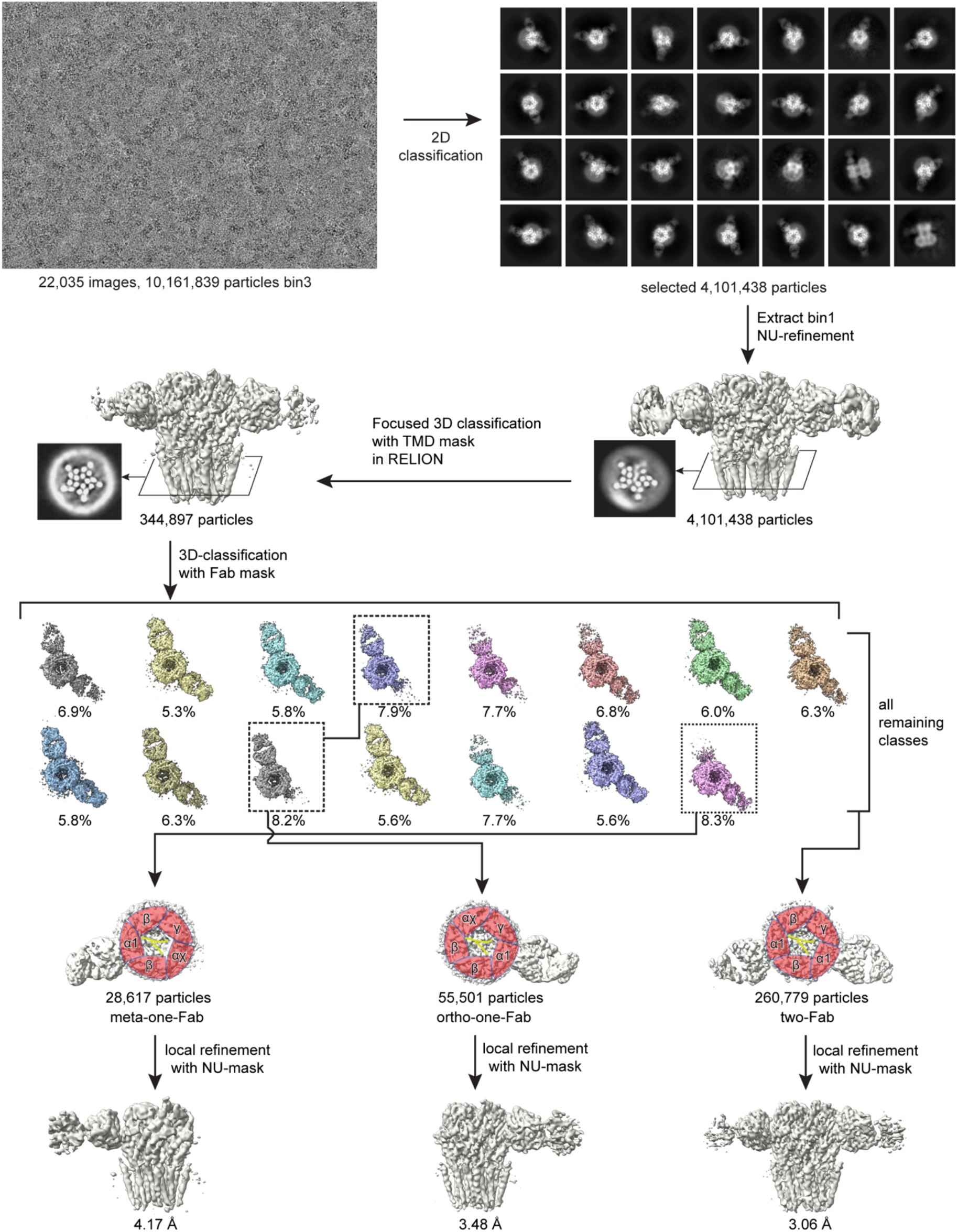
Cryo-EM data processing of the DID dataset.

**Extended Data Figure 3.**
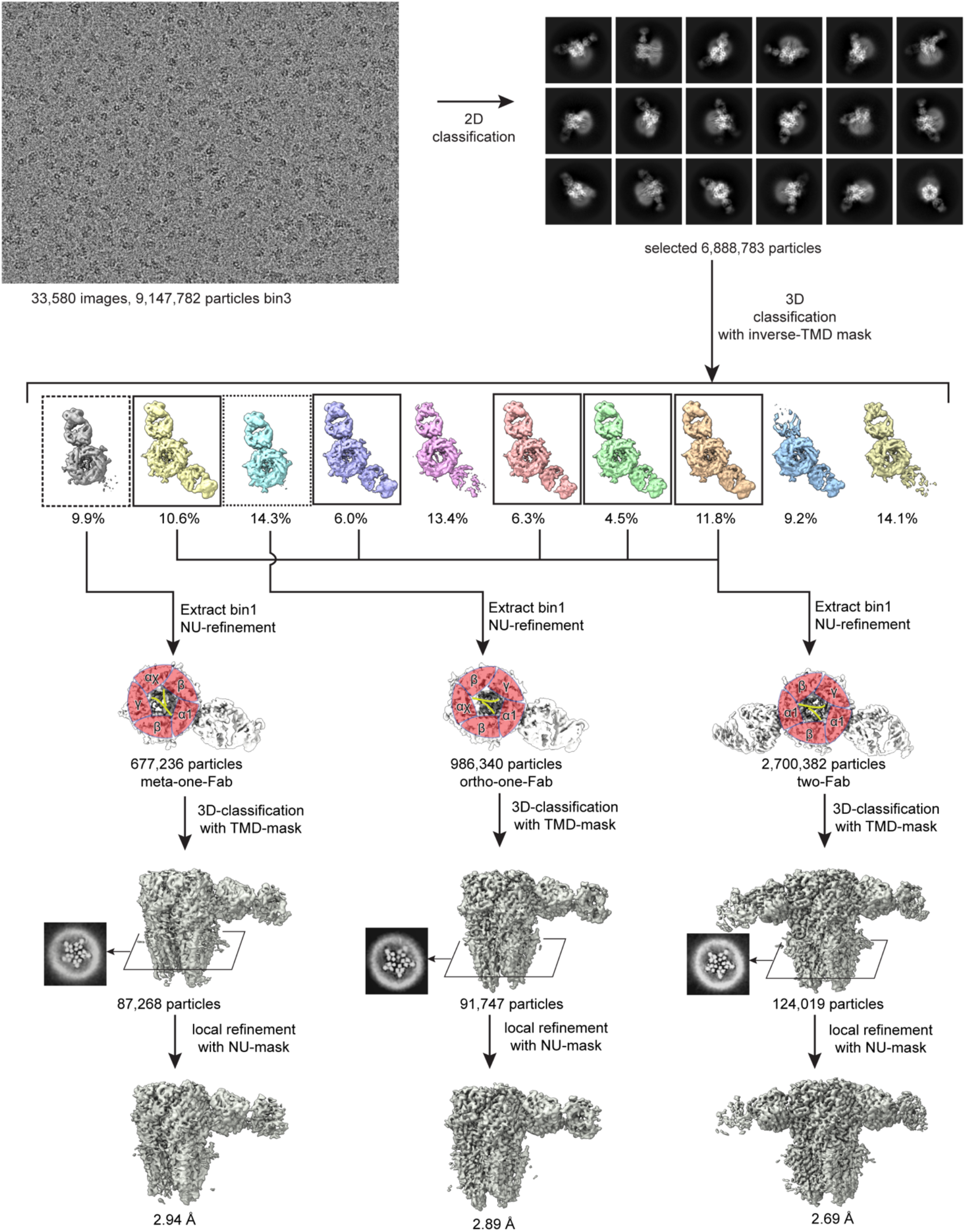
Cryo-EM data processing of the ZOL/GABA dataset.

**Extended Data Figure 4.**
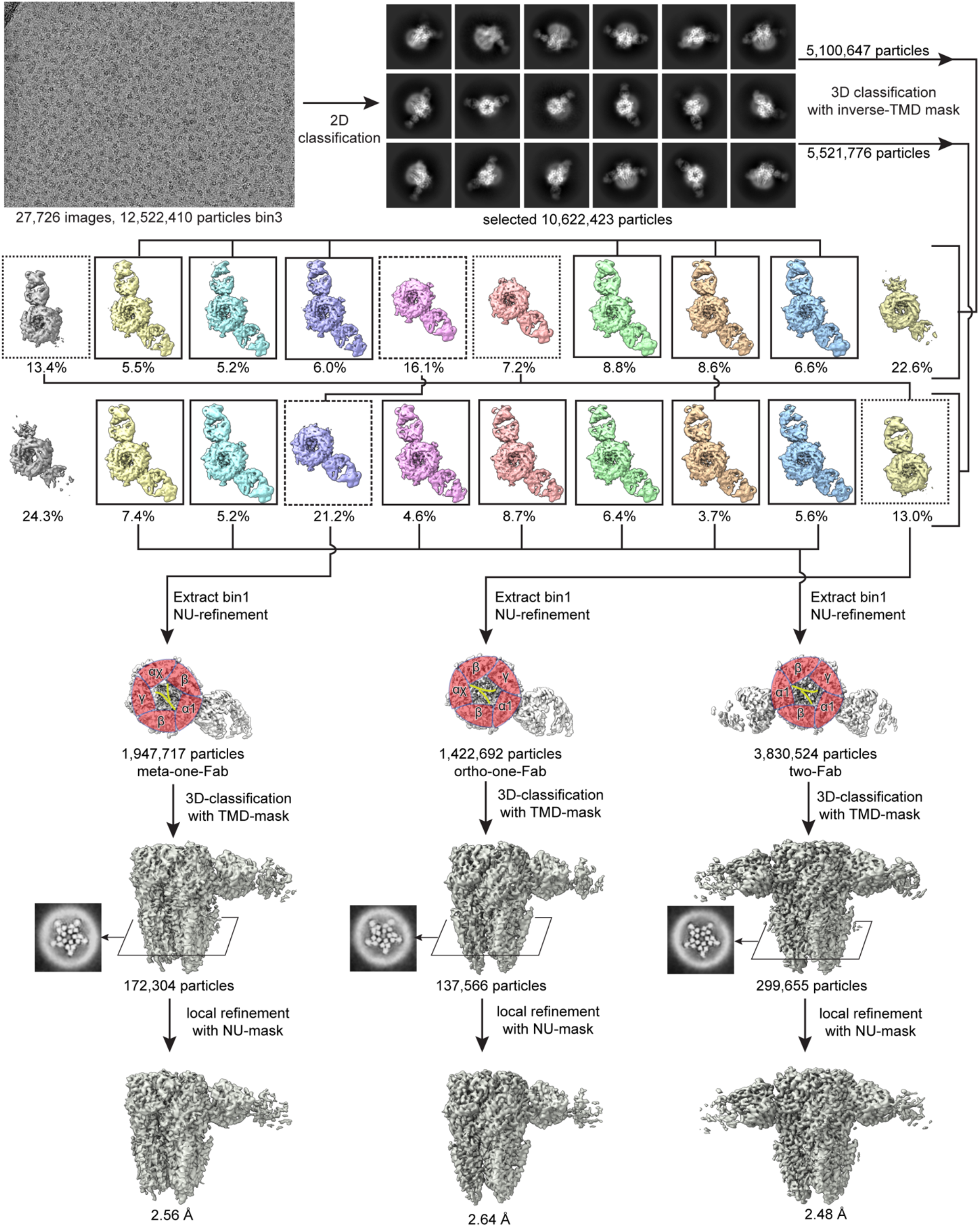
Cryo-EM data processing of the ALP/GABA dataset.

**Extended Data Figure 5.**
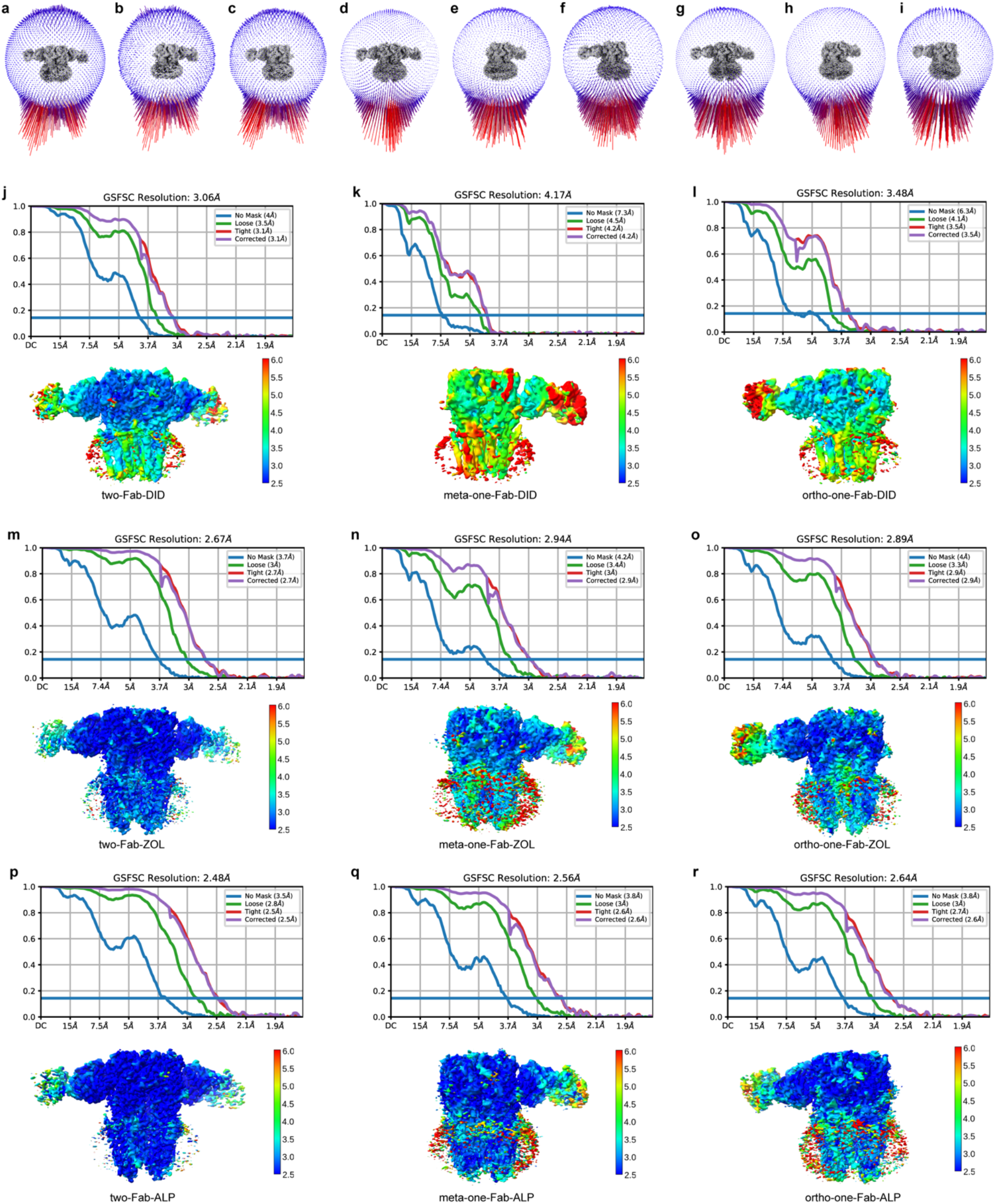
Statistics of final cryo-EM reconstructions. **a–i**, Euler angle distributions of particles used for final cryo-EM reconstruction of two-Fab-DID (**a**), meta-one-Fab-DID (**b**), ortho-one-Fab-DID (**c**), two-Fab-ZOL (**d**), meta-one-Fab-ZOL (**e**), ortho-one-Fab-ZOL (**f**), two-Fab-ALP (**g**), meta-one-Fab-ALP (**h**), ortho-one-Fab-ALP (i). **j–r**, FSC curves and local resolution plots of final cryo-EM reconstructions.

**Extended Data Figure 6.**
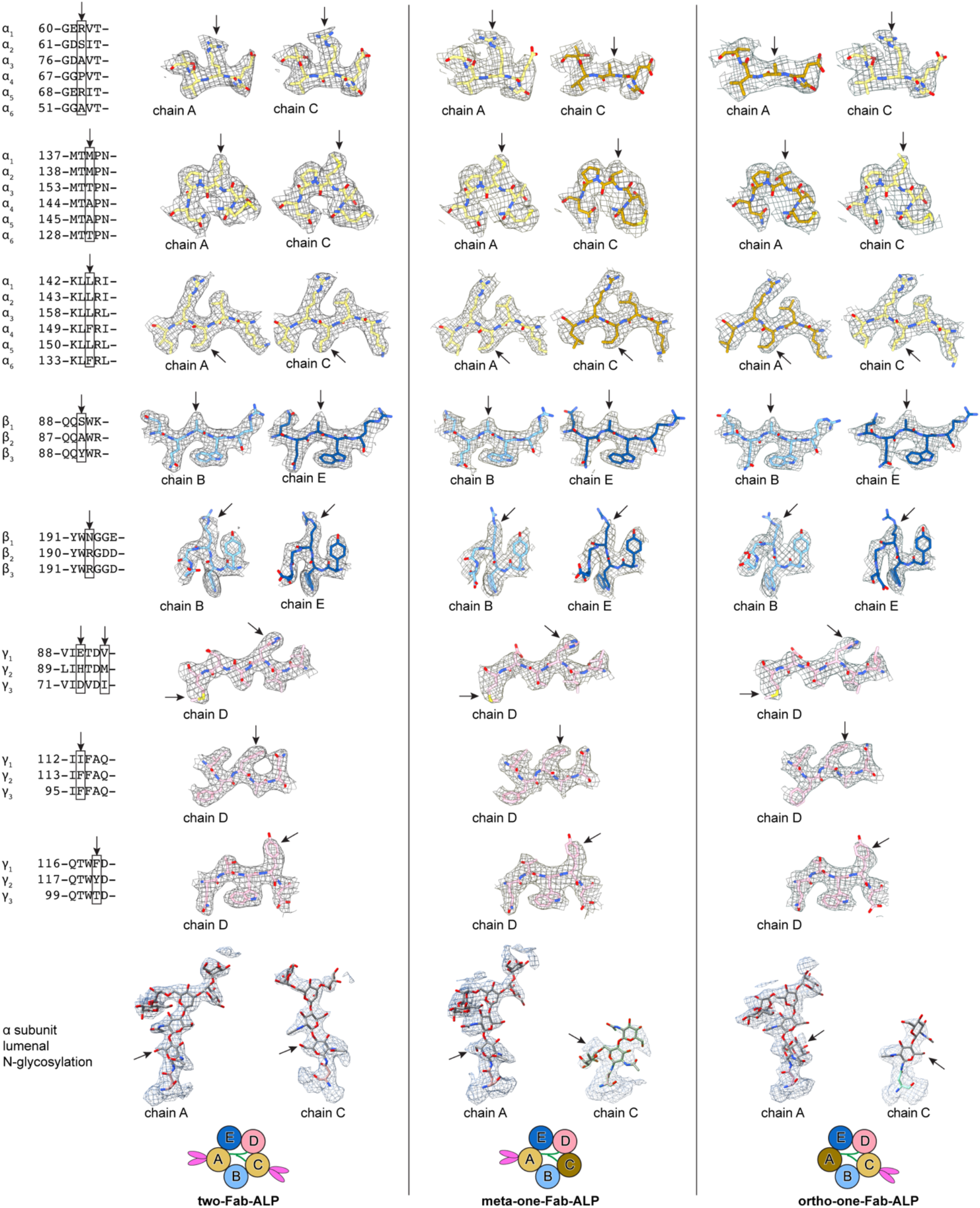
Cryo-EM densities of protein side chains and N-glycosylation used for subunit identification.

**Extended Data Figure 7.**
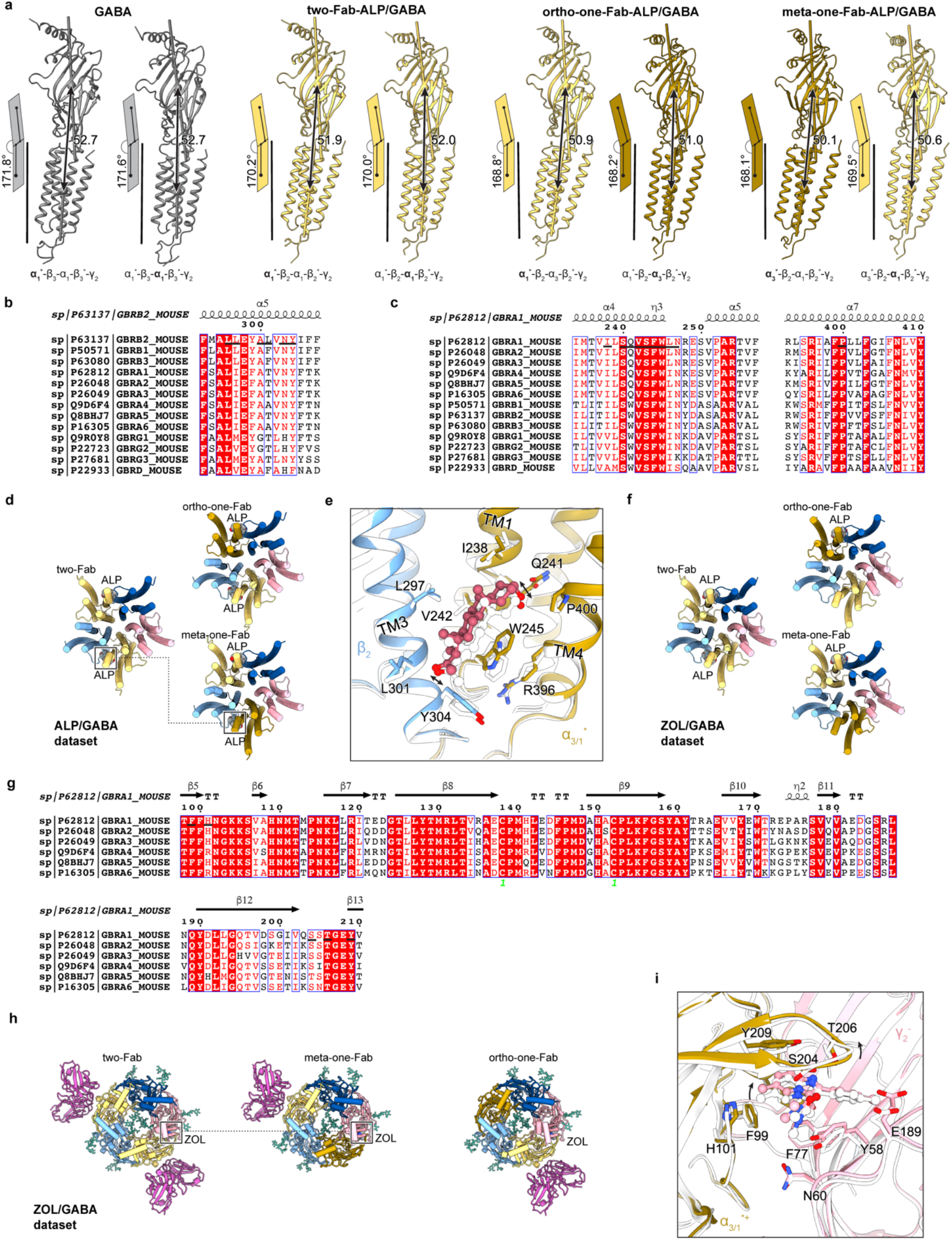
Sequence alignments of nα_1_GABA_A_R subunits and structural variations of nα_1_GABA_A_R assemblies. **a**, differential inter-domain arrangements of α subunits from the nα_1_GABA_A_Rs of the ALP/GABA dataset and a previous GABA_A_R structure (PDB code: 6I53). **b**, **c**, sequence alignments of nα_1_GABA_A_R subunits with sequence ranges relevant to neurosteroid binding. **d**, TMD structures from the ALP/GABA dataset with the allopregnanolone (ALP) shown in Vdw representation. **e**, structure comparison of the ALP binding pockets between two-Fab and meta-one-Fab. The two structures are overlayed based on the TMD of adjacent β and α subunits. **f**, TMD structures from the ZOL/GABA dataset with the endogenous neurosteroid molecules shown in Vdw representation. **g**, sequence alignments of nα_1_GABAAR α subunits with sequence ranges relevant to zolpidem binding. **h**, ECD structures from the ZOL/GABA dataset with the zolpidem shown in Vdw representation. **i**, structure comparison of the ZOL binding pockets between two-Fab and meta-one-Fab. The two structures are overlayed based on the ECD of adjacent β and α subunits.

**Extended Data Figure 8.**
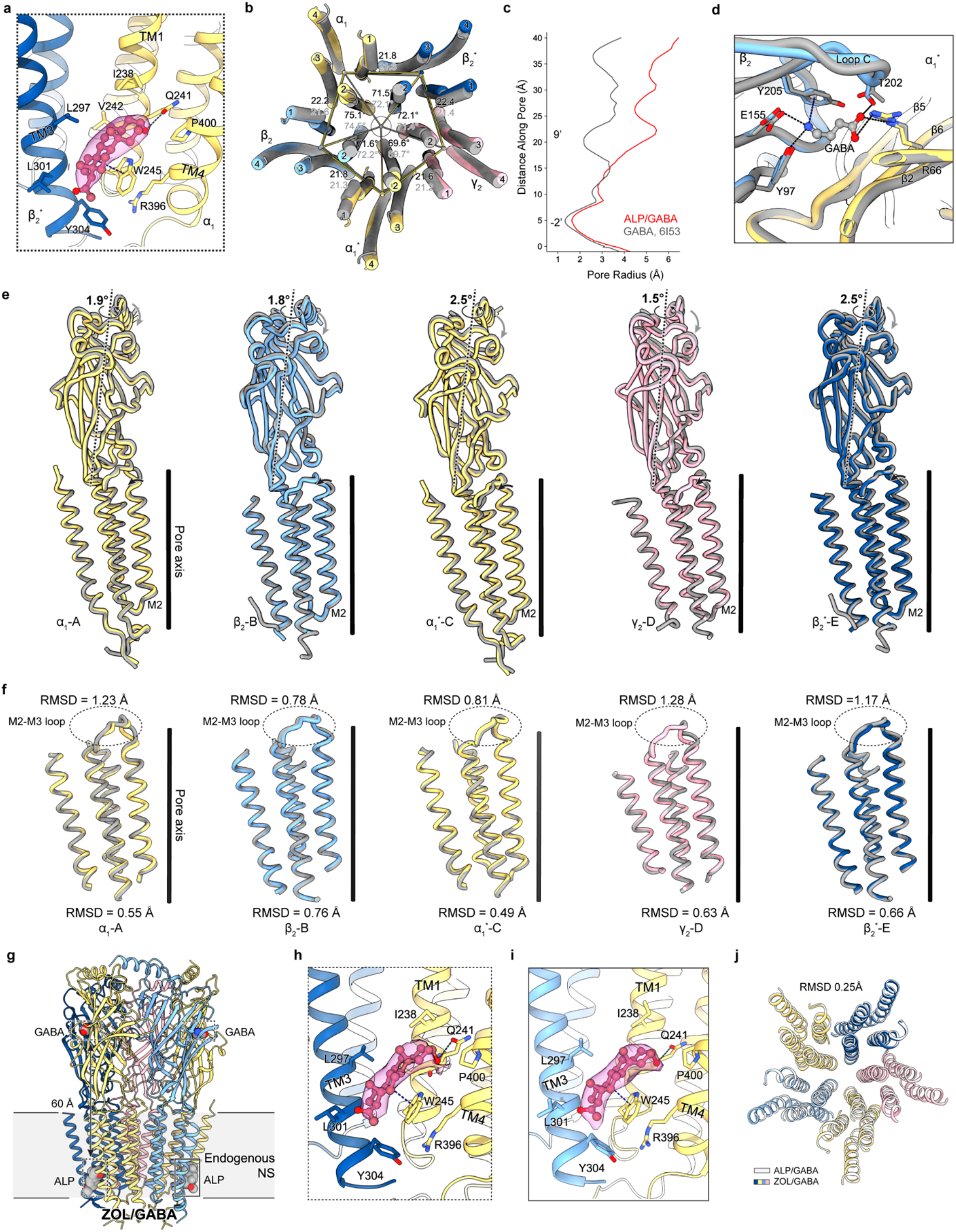
Neurosteroid binding to the nα_1_GABA_A_Rs. **a**, allopregnanolone bound between the β_2_^*+^/α_1_^-^ interface of the two-Fab-ALP. **b**, structural comparison between two-Fab-ALP and previous structure without ALP (PDB code: 6I53, apo structure hereafter). The two structures are aligned based on the global TMD. Distances and angles formed with mass centers of TMD are also shown with those of the two-Fab-ALP colored black. **c**, comparison of pore profiles between two-Fab-ALP and previous apo structure. **d**, structural overlay of the GABA binding pocket from the two-Fab-ALP and previous apo structure. The two structures are aligned based on the ECD domains of the adjacent β and α subunits. **e**, comparison of each subunit between two-Fab-ALP and previous apo structure based on global TMD structural alignment. **f**, comparison of each TMD between two-Fab-ALP and previous apo structure based on individual TMD structural alignment. RMSD values of the entire TMD domain and the M2-M3 loop are also shown. **g**, structural overview of the two-Fab-ZOL. Two ALP molecules are modeled based on the cryo-EM densities. **h**, **i**, binding poses of ALP in the two-Fab-ZOL structure. **j**, structural overlay of the two-Fab-ALP and two-Fab-ZOL based on the global TMD.

**Extended Data Figure 9.**
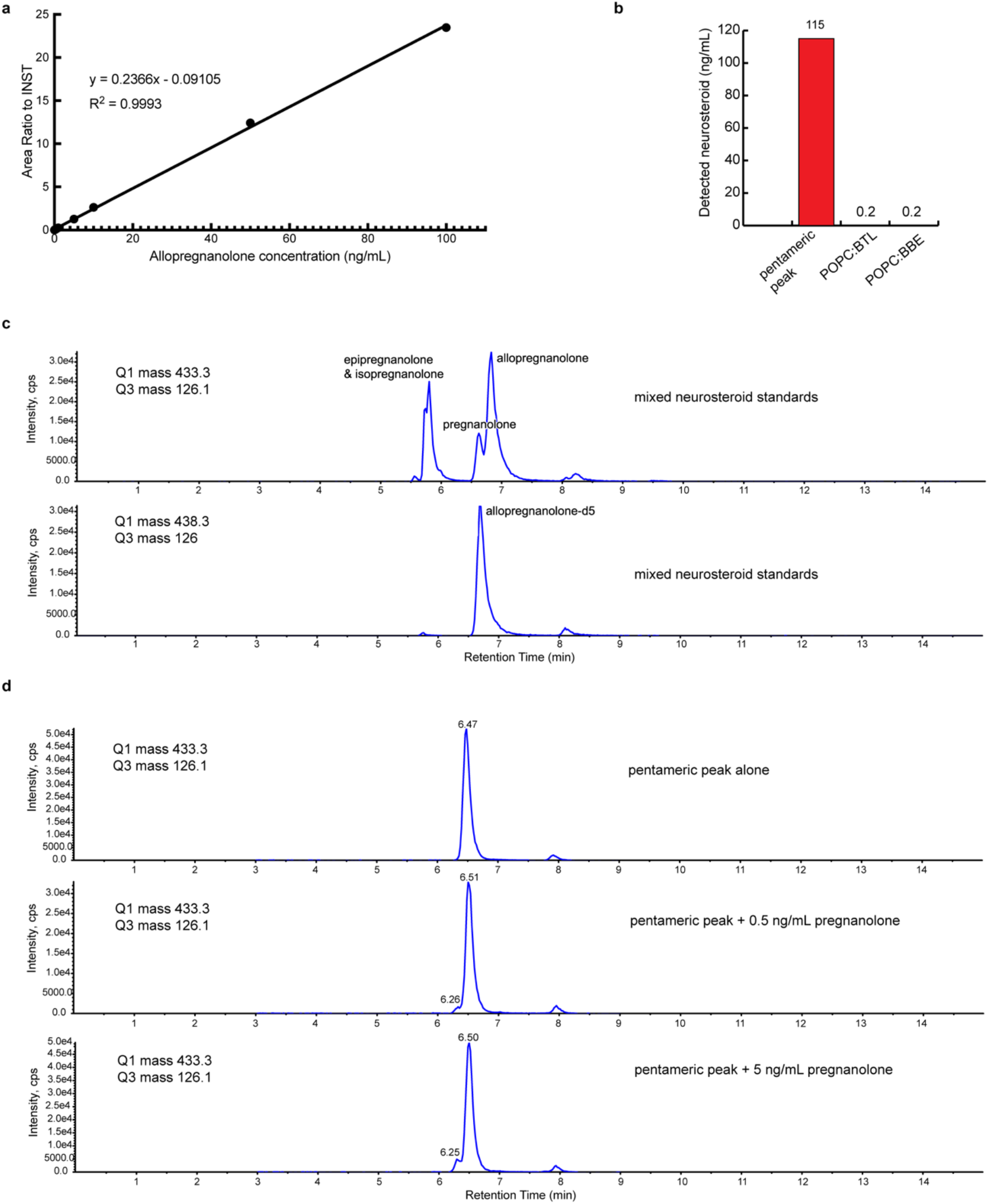
Mass spectrometry analysis of neurosteroid for the ZOL/GABA sample. **a**, standard curve of allopregnanolone quantitation based on isotope dilution. **b**, quantitation of neurosteroid in the pentameric sample and lipid stocks (85:15 mixture of POPC with either brain total lipids or bovine brain extracts) used for on-column nanodisc reconstitution. Taking into amount of the volume, the neurosteroid from the exogenous lipid makes up only 0.3% of the detected neurosteroid in the protein sample. **c**, chromatographs of the mixed neurosteroid standards under the final LC condition. **d**, chromatographs of protein sample alone or spiked with different amounts of pregnanolone standard.

**Extended Data Table 1.**
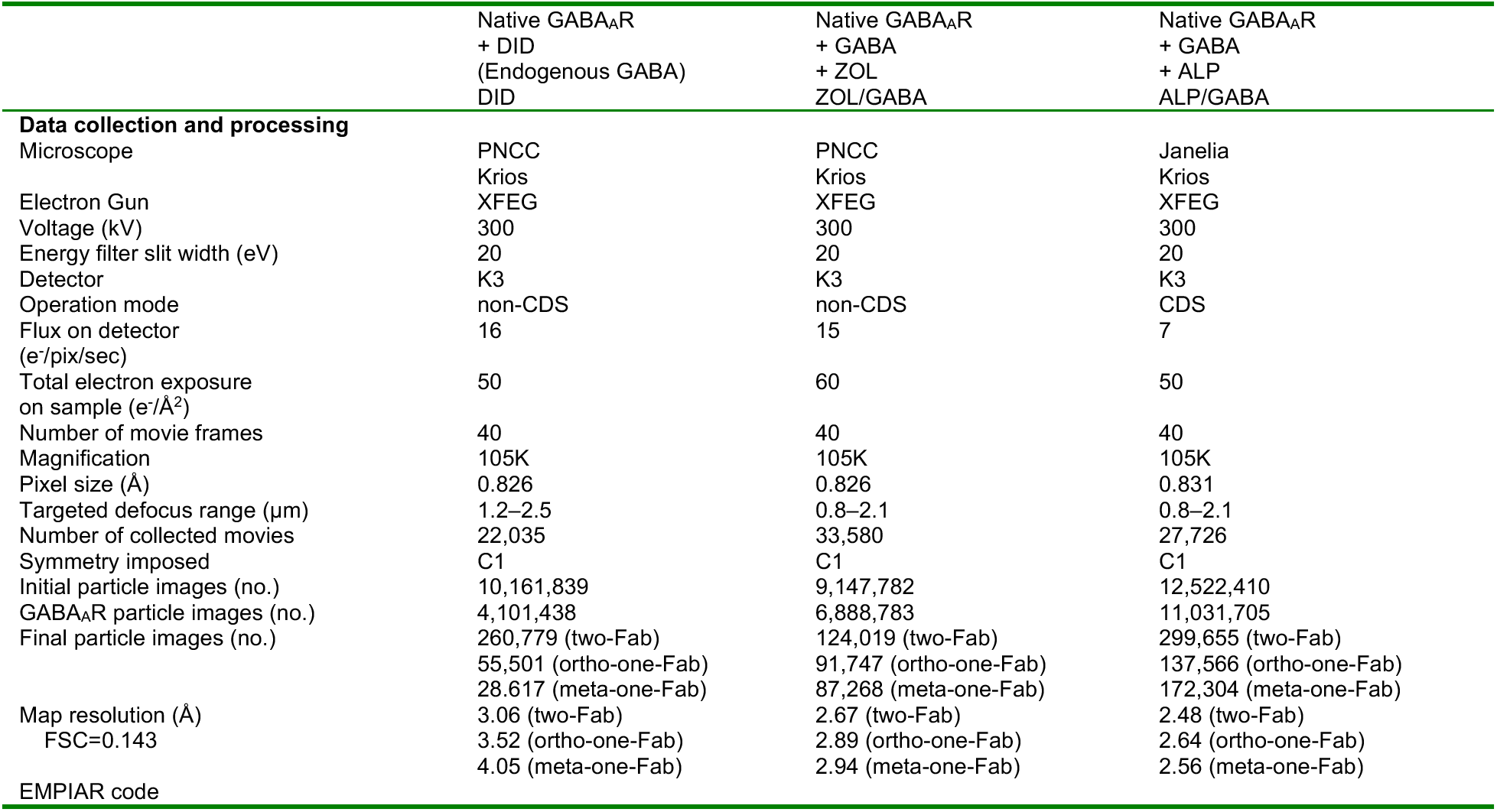
Cryo-EM data collection parameters.

**Extended Data Table 2.** Refinement and validation statistics.

|  | Two-Fab<br>+ DID<br>(Endogenous<br>GABA) | Two-Fab<br>+ GABA<br>+ ZOL<br>(Endogenous<br>neurosteroid) | Ortho-one-Fab<br>+ GABA<br>+ ZOL<br>(Endogenous<br>neurosteroid) | Meta-one-Fab<br>+ GABA<br>+ ZOL<br>(Endogenous<br>neurosteroid) | Two-Fab<br>+ GABA<br>+ ALP | Ortho-one-Fab<br>+ GABA<br>+ ALP | Meta-one-Fab<br>+ GABA<br>+ ALP |
| --- | --- | --- | --- | --- | --- | --- | --- |
| **EMDB ID** | EMD-29728 | EMD-29727 | EMD-29743 | EMD-29742 | EMD-29350 | EMD-29741 | EMD-29733 |
| **PDB ID** | 8G4O | 8G4N | 8G5H | 8G5G | 8FOI | 8G5F | 8G4X |
| Model resolution (Å) | 3.06 | 2.67 | 2.89 | 2.94 | 2.48 | 2.64 | 2.56 |
| FSC=0.143 |  |  |  |  |  |  |  |
| Map sharpening *B* factor (Å<sup>2</sup>) | 0 | 0 | 0 | 0 | 0 | 0 | 0 |
| **Model composition** |  |  |  |  |  |  |  |
| Non-hydrogen atoms | 17,355 | 18,311 | 15,876 | 15,861 | 18,288 | 15,753 | 15,899 |
| Protein residues | 2,094 | 2,120 | 1,887 | 1,887 | 2,120 | 1,888 | 1,895 |
| Glycans (molecules) | 441 (36) | 441 (36) | 430 (35) | 419 (34) | 441 (36) | 294 (23) | 386 (31) |
| Ligands (molecules) | 37 (3) | 83 (5) | 60 (4) | 60 (4) | 60 (4) | 60 (4) | 60 (4) |
| Lipids (molecules) | - | 694 (41) | 94 (2) | 94 (2) | 694 (41) | 94 (2) | 94 (2) |
| **B factors (Å<sup>2</sup>)** |  |  |  |  |  |  |  |
| Protein | 156 | 125 | 127 | 149 | 115 | 134 | 127 |
| Glycans | 212 | 171 | 179 | 204 | 165 | 170 | 179 |
| Ligands | 114 | 115 | 116 | 134 | 111 | 158 | 116 |
| Lipids | - | 144 | 182 | 205 | 146 | 199 | 172 |
| **R.m.s. deviations** |  |  |  |  |  |  |  |
| Bond lengths (Å) | 0.003 | 0.003 | 0.005 | 0.006 | 0.004 | 0.003 | 0.004 |
| Bond angles (°) | 0.526 | 0.560 | 0.999 | 0.764 | 0.627 | 0.676 | 0.683 |
| **Validation** |  |  |  |  |  |  |  |
| MolProbity score | 1.92 | 1.65 | 1.87 | 2.27 | 1.50 | 1.71 | 1.78 |
| Clashscore | 5.82 | 5.76 | 6.22 | 8.39 | 3.75 | 4.95 | 5.10 |
| Poor rotamers (%) | 3.45 | 2.24 | 2.49 | 6.23 | 1.97 | 2.37 | 2.95 |
| **Ramachandran plot** |  |  |  |  |  |  |  |
| Favored (%) | 96.81 | 97.66 | 96.51 | 96.67 | 97.47 | 97.05 | 97.17 |
| Allowed (%) | 3.19 | 2.34 | 3.49 | 3.33 | 2.53 | 2.95 | 2.83 |
| Disallowed (%) | 0.00 | 0.00 | 0.00 | 0.00 | 0.00 | 0.00 | 0.00 |

